# Phylogenomics invokes the clade housing Cryptista, Archaeplastida, and *Microheliella maris*

**DOI:** 10.1101/2021.08.29.458128

**Authors:** Euki Yazaki, Akinori Yabuki, Ayaka Imaizumi, Keitaro Kume, Tetsuo Hashimoto, Yuji Inagaki

## Abstract

As-yet-undescribed branches in the tree of eukaryotes are potentially represented by some of “orphan” protists (unicellular micro-eukaryotes), of which phylogenetic affiliations have not been clarified in previous studies. By clarifying the phylogenetic positions of orphan protists, we may fill the previous gaps in the diversity of eukaryotes and further uncover the novel affiliation between two (or more) major lineages in eukaryotes. *Microheliella maris* was originally described as a member of the phylum Heliozoa, but a pioneering large-scale phylogenetic analysis failed to place this organism within the previously described species/lineages with confidence. In this study, we analyzed a 319-gene alignment and demonstrated that *M. maris* represents a basal lineage of one of the major eukaryotic lineages, Cryptista. We here propose a new clade name “Pancryptista” for Cryptista plus *M. maris*. The 319-gene analyses also indicated that *M. maris* is a key taxon to recover the monophyly of Archaeplastida and the sister relationship between Archaeplastida and Pancryptista, which is collectively called as “CAM clade” here. Significantly, Cryptophyceae tend to be attracted to Rhodophyta depending on the taxon sampling (ex., in the absence of *M. maris* and Rhodelphidia) and the particular phylogenetic “signal” most likely hindered the stable recovery of the monophyly of Archaeplastida in previous studies. We hypothesize that many cryptophycean genes (including those in the 319-gene alignment) recombined partially with the homologous genes transferred from the red algal endosymbiont during secondary endosymbiosis and bear a faint phylogenetic affinity to the rhodophytan genes.

## 1. Introduction

Our understanding of the evolutionary relationship among major eukaryotic groups has been progressed constantly. The foundation of the tree of eukaryotes was developed initially based on the combination of morphological characteristics (including those on the ultrastructural level) and molecular phylogenetic analyses of a single or few marker genes ^1–3^. In recent years, “phylogenomic” analyses—phylogenetic analyses of large-scale multigene alignments, particularly those comprising hundreds of genes—were often conducted to reconstruct deep splits in the tree of eukaryotes with high statistical support ^4–6^. For instance, recent phylogenomic analyses have constantly reconstructed the clade of stramenopiles, Alveolata, and Rhizaria (SAR clade) ^7^, that of Opisthokonta, Amoebozoa, Breviatea, and Apusomonadida (Amorphea) ^8^, and that of Collodictyonidae, Rigifilida, and *Mantamons* (CRuMs) ^9^.

There are many unicellular micro-eukaryotes/lineages of which phylogenetic positions remain uncertain (“orphan” eukaryotes/lineages). Some of the current orphan eukaryotes/lineages most likely represent as-yet-unknown portions of the diversity of eukaryotes and hold clues to resolve the eukaryotic evolution. Prior to DNA sequencing experiments gaining in popularity in phylogenetic/taxonomic studies, diverse eukaryotes were isolated from the natural environments and examined by microscopes. If the morphological characteristics of the eukaryotes of interest showed no clear affinity to any other eukaryotes, their phylogenetic affiliations remained uncertain ^10–14^. The analyses of small subunit ribosomal DNA (SSU rDNA)—one of the most popular gene markers for organismal phylogeny—succeeded in finding the phylogenetic homes of many eukaryotes/lineages, of which morphological information was insufficient to resolve their phylogenetic affiliations ^15–18^. More recently, orphan eukaryotes/lineages, as well as newly found eukaryotes have been subjected to phylogenomic analyses ^7,8,19–26^.

Phylogenomic analyses are unlikely the silver bullet to all of the orphan eukaryotes/lineages recognized to date. For instance, the positions of Malawimonadida ^27^, Ancyromonadida ^9^, Hemimastigophora ^28^, *Ancoracysta twista* ^29^, and *Microheliella maris* ^30^ could not be clarified even after phylogenomic analyses, implying that they are genuine deep branches that are critical to resolving the backbone of the tree of eukaryotes ^9,28–30^. In this study, we challenged to clarify the phylogenetic position of *M. maris* by analyzing a new phylogenomic alignment. *M. maris* was originally described as a member of the phylum Heliozoa based on the shared morphological similarities (e.g., the radiating axopodia with tiny granules and the centroplast) ^31^. Cavalier-Smith et al. (2015) ^30^ then examined the phylogenetic position of *M. maris* by analyzing the alignment comprising 187 genes. Nevertheless, *M. maris* is still regarded as one of the orphan eukaryotes ^32^, as the choice of the methods for tree reconstruction and taxon sampling affected largely the position of this eukaryote in the 187-gene phylogeny ^30^.

We here reassessed the phylogenetic position of *M. maris* by analyzing a new phylogenomic alignment comprising 319 genes (88,592 amino acid positions in total). The 319-gene phylogeny placed *M. maris* at the base of the Cryptista clade with high statistical support, suggesting that this eukaryote holds keys to understand the early evolution of Cryptista as well as Diaphoretickes. Indeed, we further demonstrated that *M. maris* and Rhodelphidia, which occupy the basal position of Cryptista and that of Rhodophyta, respectively, suppress the erroneous ‘signal’ attracting Cryptophyceae and Rhodophyta to each other and contribute to recovering (i) the monophyly of Archaeplastida and (ii) the sister relationship between Archaeplastida and the clade of Cryptista plus *M. maris*. Finally, we explored the biological ground for the phylogenetic artifact uniting Cryptophyceae and Rhodophyta together.

## 2. Methods

### (a) Cell culturing and RNA-seq analysis

We generated the RNA-seq data from *M. maris* and *Hemiarma marina*, a species of Goniomonadea, in this study. The culture of *M. maris* [studied in Yabuki et al. (2012) ^31^] and that of *H. marina* [established in Shiratori and Ishida (2016) ^33^] have been kept in the laboratory and were utilized in this study. The harvested cells of both organisms were subjected to RNA extraction using TRIzol (Life Technologies) by following the manufacturer’s instructions. We shipped the two RNA samples to a biotech company (Hokkaido System Science) for cDNA library construction from the poly-A-tailed RNAs followed by sequencing using the Illumina Hi-seq 2000 platform. For *M. maris*, 1.6 × 10^7^ paired-end 100 bp reads (1.6 Gb in total) were obtained and then assembled into 30,305 unique contigs by TRINITY ^34,35^. For *H. marina*, we obtained 1.9 × 10^7^ paired-end 100 bp reads (1.9 Gb in total) and assembled them into 41,539 unique contigs by TRINITY ^34,35^.

### (b) Global eukaryotic phylogeny

To elucidate the phylogenetic position of *M. maris*, we prepared a phylogenomic alignment by updating an existing dataset comprising 351 genes ^28^. For each of the 351 genes, we added the homologous sequences retrieved from the transcriptomic data newly generated from *M. maris* and *H. marina* in this study (see above), as well as other eukaryotes that were absent in the original data ^28^, such as *Marophrys* sp. SRT127 ^36^, two species of Rhodelphidia (i.e., *Rhodelphis limneticus* and *R. marinus*) ^25^, and *Ancoracysta twista* ^29^. Individual single-gene alignments were aligned by MAFFT v. 7.205 ^37,38^ with the L-INS-i algorithm followed by manual correction and exclusion of ambiguously aligned positions. For each of the single-gene alignments, the maximum-likelihood (ML) phylogenetic tree was inferred by FASTTREE v. 2.1 ^39,40^ under the LG + Γ model with robustness assessed with a 100-replicate bootstrap analysis. Individual single-gene trees were inspected to identify the alignments bearing aberrant phylogenetic signal that disagreed strongly with any of a set of well-established monophyletic assemblages in the tree of eukaryotes, namely Opisthokonta, Amoebozoa, Alveolata, stramenopiles, Rhizaria, Rhodophyta, Chloroplastida, Glaucophyta, Haptophyta, Cryptista, Jakobida, Euglenozoa, Heterolobosea, Diplomonadida, Parabasalia, and Malawimonadida. 32 out of the 351 single-gene alignments were found to violate the above-mentioned criteria and were excluded from the phylogenomic analyses described below. The remaining 319 single-gene alignments (Table S1) were concatenated into a single phylogenomic alignment containing 82 taxa with 88,592 unambiguously aligned amino acid positions. The coverage for each single-gene alignment is summarized in Table S1.

We first subjected the final alignment comprising 319 genes from 82 taxa (GlobE alignment) to the ML method by IQ-TREE 1.6.12 ^41^ with the LG + Γ + F model, and the resultant ML tree was then used as a guide tree in a more thorough analysis with the LG + Γ + F + C60 + PMSF (posterior mean site frequencies) model ^42^. The robustness of the ML phylogenetic tree was evaluated with a nonparametric ML bootstrap analysis with the LG + Γ + F + C20 + PMSF model (100 replicates). We also conducted Bayesian phylogenetic analysis with the CAT + GTR model using PHYLOBAYES-mpi v. 1.8a ^43,44^. In this analysis, two MCMC runs were run for 5,000 cycles with ‘burn-in’ of 1,250. The consensus tree with branch lengths and Bayesian posterior probabilities (BPPs) were calculated from the remaining trees.

We evaluated the contribution of fast-evolving positions in the GlobE alignment to the position of *M. maris*. Substitution rates of individual alignment positions were calculated over the ML tree by IQ-TREE 1.6.12 ^41^ and top 20%, 40%, 60%, and 80% fastest-evolving positions were then removed from the original alignment. The processed alignments were then subjected to the ML bootstrap analysis with the UFBOOT approximation ^45^ (1,000 replicates) by using IQ-TREE 1.6.12 ^41^ with the LG + Γ + F model. Henceforth, the alignment modification and the following ML bootstrap analyses are designated as “FPR (fast-evolving position removal)” analysis.

We also examined the impact of the sampling of the genes in the GlobE alignment on the position of *M. maris* by “RGS (random gene sampling)” analyses described below ^46^. From the 319 genes in the GlobE alignment, 50 genes were randomly sampled and concatenated into a single alignment (“rs50g” alignment). The above procedure was repeated 50 times to obtain 50 of rs50g alignments. Likewise, we prepared (i) 50 of “rs100g” alignments comprising 100 randomly sampled genes, (ii) 10 of “rs150g” alignments comprising 150 randomly sampled genes, and (iii) 10 of “rs200g” alignments comprising 200 randomly sampled genes. The alignments comprising randomly sampled genes were subjected individually to the ML bootstrap analysis with the UFBOOT approximation (1,000 replicates) by using IQ-TREE 1.6.12 ^41^ with the LG + Γ + F model.

### (c) Diaphoretickes phylogeny

To evaluate the impact of the inclusion of *M. maris* to the phylogenetic relationship among the species/lineages in Dipahoretickes, we excluded 22 taxa from the GlobE alignment to generate the second phylogenomic alignment, of which taxa were occupied mostly by members of Diaphoretickes. Note that the number of genes remained the same between the GlobE and the second, “Diaph” alignments. The Diaph alignment was subjected to both ML and Bayesian phylogenetic analysis under all the same conditions as described above, except that we used the LG + Γ + F + C60 + PMSF model for the ML bootstrap analysis. Both FPR and RGS analyses (see above) were applied to the Diaph alignment. We also conducted both FPR and RGS analyses after excluding Rhodelphidia and *M. maris* alternatively from the Diaph alignment.

The taxon sampling of the Diaph alignment was further modified by excluding (i) both Rhodelphidia and *M. maris*, (ii) Rhodelphidia, *M. maris*, and *P. bilix*, (iii) Rhodelphidia, *M. maris*, *P. bilix*, and Goniomonadea, and (iv) Rhodelphidia, *M. maris*, *P. bilix*, and Cryptophyceae. We ran RGS analyses of all of the four alignments described above, and the last two were subjected to FPR analyses as well.

## 3. Results and Discussion

### (a) *Microheliella maris* represents a lineage basal to Cryptista: Proposal of “Pancryptista”

We analyzed a transcriptome-based GlobE alignment consisting of 319 genes sampled from 82 eukaryotes, which represent the major taxonomic assemblages and several orphan taxa/lineages. The GlobE phylogeny recovered the major clades in eukaryotes, such as SAR, Amorphea, CRuMs, Discoba, and Cryptista with full statistical support in both ML and Bayesian methods (Fig. 1; see also Fig. S1). *M. maris* branched at the base of the Cryptista clade, which comprises *Palpitomonas bilix*, Goniomonadea including *Hemiarma marina*, and Cryptophyceae, with an MLBP of 100% and a BPP of 1.0. The GlobE alignment includes no data of Kathablepharidacea that is the other cryptistan subgroup, as their available data is extremely low site coverage. However, the lack of Kathablepharidacea most likely gave little impact on the phylogenetic position of *M. maris* relative to Cryptista, as far as the alignment includes *P. bilix* which is more basal than Kathablepharidacea in the Cryptista clade ^20^. The monophyly of Archaeplastida including Rhodelphidia (*Rhodelphis limneticus* and *R. marinus*) and Picozoa sp., both of which grouped with Rhodophyta, was recovered with an MLBP of 81% and a BPP of 1.0. The intimate affinity of Rhdelphidia and Picozoa to Rhodophyta in the GlobE phylogeny is consistent with the recent phylogenomic studies ^25,47^. Neither of the two recently proposed major clades in eukaryotes, T-SAR (Telonemia plus SAR) ^23^ and Haptista (Centrohelea plus Haptophyta) ^21^, was reconstructed. Either or both ML and Bayesian phylogenetic analyses failed to give full statistical support to the nodes connecting the lineages/species in Diaphoretickes, namely Archaeplastida, Centrohelea, Haptophyta, Telonemia, SAR, and Cryptista plus *M. maris*. Thus, we conclude that the 82-taxon phylogeny is insufficient to retrace the early evolution of Diaphoretickes with confidence.

**Figure 1.**
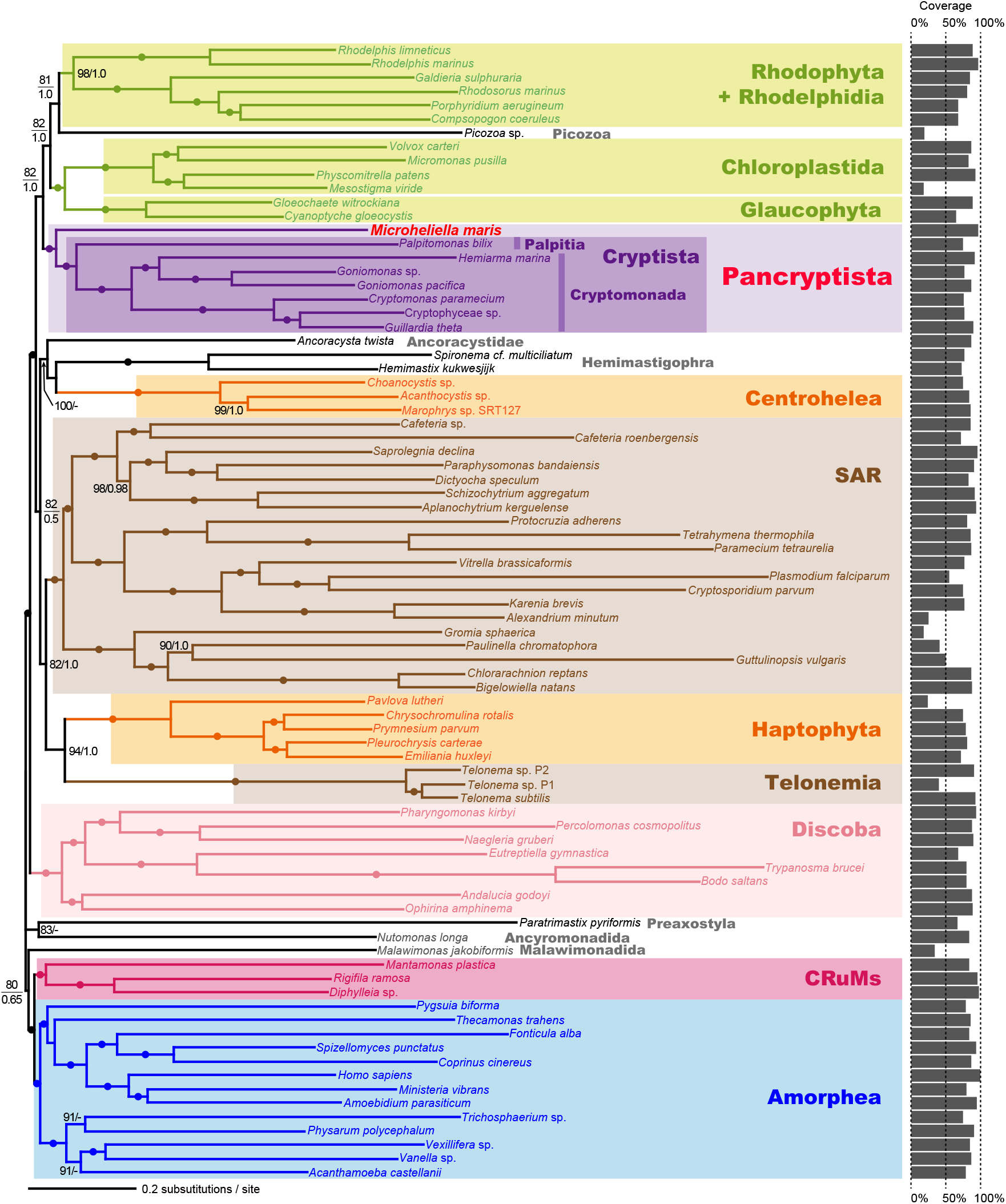
Phylogenetic position of *Microheliella maris* inferred from the GlobE alignment. The tree topology and branch lengths were inferred from the GlobE alignment (319 genes; 88,592 amino acid positions in total) by the maximum-likelihood (ML) and Bayesian methods. Bayesian analysis recovered principally an identical tree topology (Fig. S1). For each bipartition, the ML bootstrap support values (MLBPs) and Bayesian posterior probabilities (BPPs; if greater than 0.50) are shown. The bipartitions with dots indicate MLBPs of 100% and BPPs of 1.0. The bar graph for each taxon represents the percent coverage of the amino acid positions in the GlobE alignment.

We here examined the phylogenetic position of *M. maris* inferred from the GlobE alignment by the progressive removal of fast-evolving positions (FPR analyses). The contribution of fast-evolving positions in the GlobE alignment to the union of *M. maris* and Cryptista is most likely negligible, as the ultrafast bootstrap support values (UFBPs) for the clade comprising *M. maris* and Cryptista stayed 100% until the top 80% fastest-evolving positions were removed (purple line in Fig. S2A). We detected two conflicting phylogenetic signals regarding the position of *M. maris* relative to the members of Cryptista included in the GlobE alignment, one placing *M. maris* at the base of the Cryptista clade and the other uniting *M. maris* and *P. bilix* directly (red and yellow lines, respectively, in Fig. S2A). However, the former signal constantly dominated over the latter, regardless of the amount of fast-evolving positions in the alignment. Thus, we conclude that the basal position of *M. maris* to the Cryptista clade in the GlobE phylogeny (Fig. 1) is free from potential phylogenetic artifacts stemming from fast-evolving positions.

To evaluate the impact of gene sampling on the position of *M. maris* in the GlobE phylogeny, we randomly sampled 50 genes, 100 genes, 150 genes, and 200 genes from the 319 genes and concatenated them into “rs50g,” “rs100g,” “rs150g,” and “rs200g” alignments, respectively (note that the taxon sampling remained the same). The UFBPs for the clade comprising *M. maris* and Cryptista calculated from 50 of rs50g alignments and 50 of rs100g alignments distributed from 0 (or nearly 0) to 100% (Fig. S2B; see also Table S2 for the details), suggesting the phylogenetic signal in 100 or fewer genes are insufficient for stable recovery of the grouping of interest. Nevertheless, in the analyses of rs150g and rs200g alignments, the UFBP for the grouping of *M. maris* and Cryptista increased. Significantly, the support for the grouping of *M. maris* and Cryptista received UFBPs of 81.3-100% in the analyses of rs200g alignments (the rightmost plot in Fig. S2B; see also Table S2). As observed in FPR analysis of GlobE alignment (Fig. S2A), rs50g and rs100g alignments appeared to contain two conflicting signals for the phylogenetic position of *M. maris*, one placing *M. maris* at the base of the Cryptista clade and the other uniting *M. maris* and *P. bilix* directly (Figs. S2C and D). In the analyses of rs150g alignments, the UFBP for the basal position of *M. maris* to the Cryptista clade appeared to be greater than that for the direct union of *M. maris* and *P. bilix* in 8 out of the 10 cases (Table S2). The same trend was observed in the analyses of rs200g alignments, albeit the signal for the clade grouping *M. maris* and *P. bilix* remained detectable (Fig. S2D). This series of the analyses demonstrated that the greater the number of genes included in the alignment, the greater the UFBP for the basal position of *M. maris* at the Cryptista clade. Based on the results described above, we conclude that the basal position of *M. maris* to the Cryptista clade is genuine, and henceforth designate the clade grouping *M. maris* and the previously known Cryptista as Pancryptista.

### (b) The sister relationship between Archaeplastida and Pancryptista: Proposal of “CAM clade”

The Diaph alignment, which was generated by excluding 22 taxa from the GlobE alignment, was analyzed to explore the impact of *M. maris* on the phylogenetic relationship among the major lineages in Diaphoretickes. 21 out of the 22 taxa excluded from the GlobE alignment were not the member of Diaphoretickes—most of Opisthokonta, all discobids, *Paratrimastix pyriformis*, *Nutomonas longa*, and *Malawimonas jakobiformis*. We excluded a single member of Diaphoretickes, Picozoa sp., from the Diaph alignment due to its instability in the GlobE phylogeny, which likely stemmed from low site coverage in the GlobE alignment (Fig. 1). The phylogenetic relationship among the major Diaphoretickes lineages/species inferred from the Diaph alignment (e.g., the basal position of *M. maris* to the Cryptista clade) was essentially the same as that inferred from the GlobE alignment (Figs. 1 and 2A). We here focus on the monophyly of Archaeplastida and the sister relationship between Archaeplastida and Pancryptista, both of which were fully supported in the ML and Bayesian analyses of the Diaph alignment (Fig. 2A).

**Figure 2.**
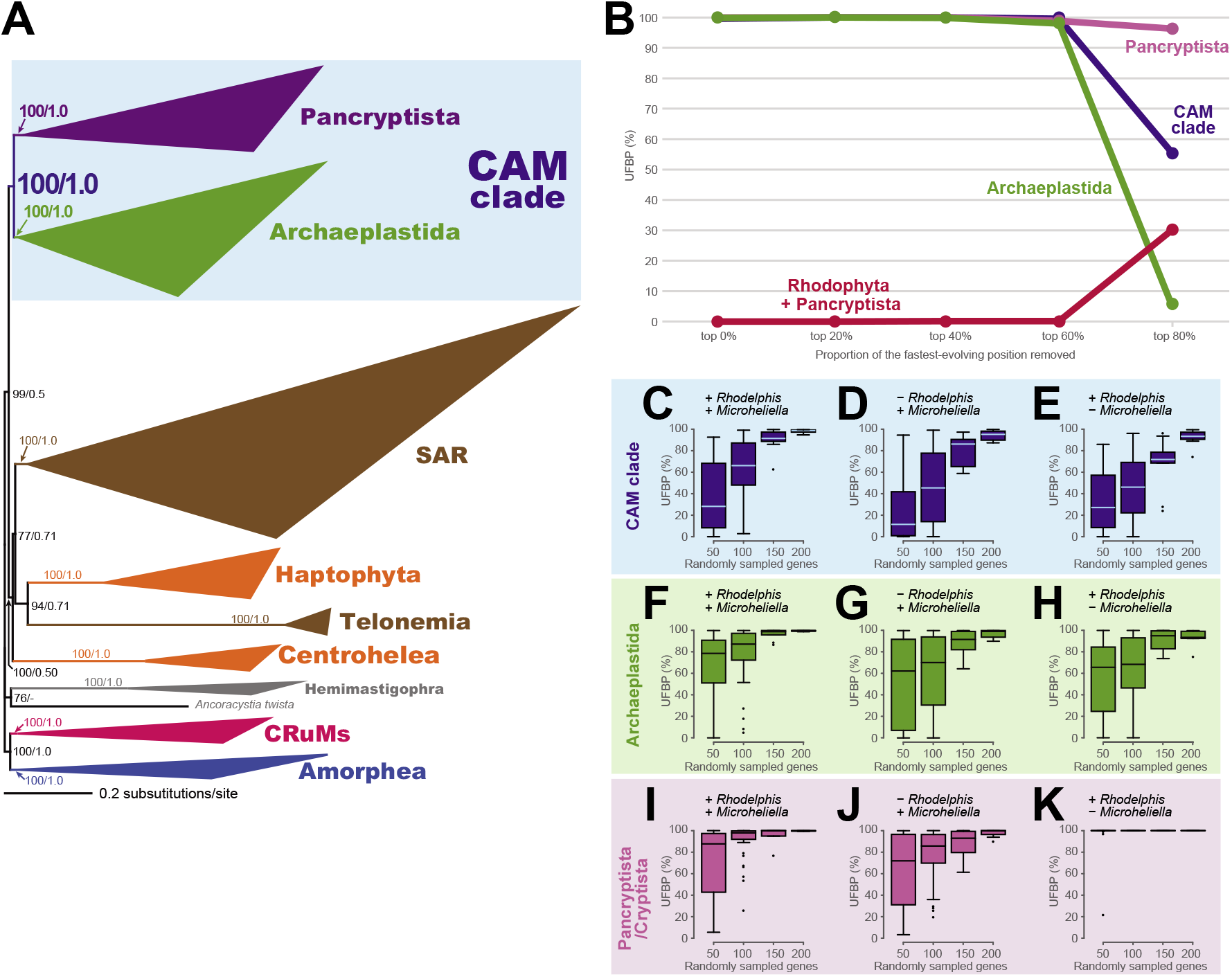
Analyses assessing the impact of *Microheliella maris* and Rhodelphidia on the monophyly of Pancryptista, the monophyly of Archaeplastida, and CAM clade. Note: We regard the sister relationship between Pancryptista and Archaeplastida as CAM clade. If Pancryptista (or Cryptista) was placed within Archaeplastida, we do not consider such assemblage as CAM clade. **(A)** The maximum-likelihood (ML) tree inferred from the Diaph alignment comprising 319 genes (88,592 amino acid positions in total). Clades of closely related taxa are collapsed as triangles. For the detailed ML tree, please refer to Fig. S2. Bayesian analysis recovered principally an identical tree topology (Fig. S3). ML bootstrap support values (MLBPs) and Bayesian posterior probabilities (BPPs; if greater than 0.50) are indicated on the bipartitions presented in the figure. **(B)** Analyses of Diaph alignment processed by fast-evolving position removal (FPR). We repeated ultrafast bootstrap analyses using IQ-TREE 1.6.12 on the Diaph alignment after excluding no position, the top 20, 40, 60, and 80% fastest-evolving positions. The plots in purple, blue, green, and red indicate the ultrafast bootstrap support values (UFBPs) for the monophyly of Pancryptista, the monophyly of Archaeplastida, CAM clade, and the union of Rhodophyta and Pancryptista, respectively. **(C-K)** Analyses of the alignments generated by random gene sampling (RGS). We sampled 50, 100, 150, and 200 genes randomly from the 319 genes in the Diaph alignment, concatenated into “rs50g,” “rs100g,” “rs150g,” and “rs200g” alignments, and subjected to ultrafast bootstrap analyses using IQ-TREE 1.6.12. We presented the UFBPs for CAM clade, the monophyly of Archaeplastida, and the monophyly of Pancryptista as box-and-whisker plots **(C)**, **(F)**, and **(I)**, respectively. The above-mentioned analyses were repeated after *Rhodelphis* spp. or *M. maris* were excluded from the alignments alternatively. The UFBPs from the analyses excluding *Rhodelphis* spp. and those from the analyses excluding *M. maris* are presented in **(D)**, **(G)**, and **(J),** and **(E)**, **(H)**, and **(K)**, respectively. The UFBPs shown in the plots described above are summarized in Table S3.

Neither monophyly of Archaeplastida nor sister relationship between Archaeplastida and Cryptista has not been settled even by recently published phylogenomic studies. For instance, Gawryluk et al. (2019) ^25^ conducted the analyses of an alignment comprising 254 genes but could not settle the relationship among Chloroplastida, Glaucophyta, and Rhodophyta plus Rhodelphidia. The ML analysis of their 254-gene alignment put Cryptista within the three lineages of Archaeplastida, albeit Bayesian analysis of the same alignment recovered both monophyly of Archaeplastida and sister relationship between Archaeplastida and Cryptista. In another phylogenomic study, the monophyly of Archaeplastida was not reconstructed in either ML or Bayesian analysis of an alignment comprising 248 genes, as Cryptista was tied with Rhodophyta ^23^. Irisarri, Strassert, and Burki (2021) ^48^ recently analyzed a 311-genes alignment and demonstrated that taxon sampling, selection of alignment positions, and substitution models for phylogeny are critical to recovering the monophyly of Archaeplastida.

In contrast to the pioneering phylogenomic studies (see above), both monophyly of Archaeplastida and the sister relationship between Pancryptista and Archaeplastida were reconstructed from the Diaph alignment with full statistical support by both ML and Bayesian methods (Fig. 2A). Significantly, FPR analysis on the Diaph alignment appeared to have little impact on the UFBPs for the two nodes of interest (Fig. 2B). Both monophyly of Archaeplastida and sister relationship between Pancryptista and Archaeplastida received full or nearly full UFBPs until the top 60% fastest-evolving positions were removed (purple and green lines, respectively, in Fig. 2B). We also analyzed rs50g, rs100g, rs150g, and rs200g alignments generated from the Diaph alignment (Figs. 2C, F, and I; see also Table S3 for the details). The results from RGS analyses clearly indicate that the UFBPs for the two nodes of interest (and that for the monophyly of Pancryptista) increased in proportion to the number of genes considered. We here conclude that Archaeplastida is a genuine clade as demonstrated by Irisarri, Strassert, and Burki (2021) ^48^, and Pancryptista is the closest relative of Archaeplastida and thus propose the sister relationship between Archaeplastida and Pancryptista as “CAM” clade. The proposed name is an acronym derived from the first letters of Cryptista, Archaeplastida, and *Microheliella*.

It is significant to note that the monophyly of Archaeplastida and the sister relationship between Archaeplastida and Cryptista was recovered by a 311-genes phylogeny prior to the *M. maris* data is available ^48^. Thus, we decided to evaluate systematically how the increment of *M. maris*, as well as that of Rhodelphidia, contributed to the recovery of the monophyly of Archaeplastida and the sister relationship between Archaeplastida and Pancryptista/Cryptista. We re-analyzed rs50g, rs100g, rs150g, and rs200g alignments generated from the Diaph alignment after excluding Rhodelphidia or *M. maris* (Figs. 2D, E, G, H, J, and K; see also Table S4 for the details). In the analyses of rs50g and rs100g alignments, the removal of Rhodelphidia/*M. maris* lowered the overall distributions of the UFBPs for the monophyly of Archaeplastida and sister relationship between Archaeplastida and Pancryptista/Cryptista (Figs. 2D, E, G, and H). However, most of the UFBPs for the two groupings of interest in the analyses of rs200g alignments were around of or greater than 90% (Figs. 2D, E, G, and H; Table S4). Likewise, the overall distribution of the UFBPs for the monophyly of Pancryptista was apparently lowered in the analyses of rs50g, rs100g, and rs150g alignments in the absence of Rhodelphidia (Fig. 2J). After *M. maris* was excluded, the monophyly of Cyrptista was constantly recovered with full UFBPs, except a UFBP of 21.5% obtained in the analyses of a single rs50g alignment (Fig. 2K; Table S4). The removal of Rhodelphidia or *M. maris* appeared to possess a moderate but apparent impact on the monophyly of Archaeplastida and sister relationship between Archaeplastida and Pancryptista/Cryptista in the alignments comprising 100 or fewer genes, albeit such impact can be overcome by an increment of the alignment size.

### (c) On the artifactual grouping of Rhodophyta and Cryptophyceae

The phylogenetic analyses described above indicated that taxon sampling is a key to recover the monophyly of Archaeplastida and the sister relationship between Archaeplastida and Pancryptista (CAM clade) with confidence. Then, why did phylogenomic analyses, in which either or both of Rhodelphidia and *M. maris* were absent, often failed to recover the monophyly of Archaeplastida? For instance, a recent phylogenomic study ^23^, which considered neither Rhodelphidia nor *M. maris*, grouped Rhodophyta and Cryptista together instead of recovering the monophyly of Archaeplastida.

The absence of both Rhodelphidia and *M. maris* gave a greater impact on the UFBP for the monophyly of Archaeplastida and that for the sister relationship between Archaeplastida and Cryptista than the absence of either of the two lineages/species. Regardless of the number of randomly sampled genes, the distributions of the UFBPs for the two groupings of interest tend to be lower than the corresponding values calculated from the analyses excluding either Rhodelphidia or *M. maris* (compare Figs. 2D and E with Fig. 3A, and Fig. 2G and H with Fig. 3E; see also Table S3 for the details). Interestingly, the faint affinity between Rhodophyta and Cryptista became detectable in the absence of Rhodelphidia and *M. maris* (Fig. 3I). After *P. bilix* was additionally excluded (Cryptista was represented by Goniomonadea and Cryptophyceae), both UFBP for the monophyly of Archaeplastida and that for the sister relationship between Archaeplastida and Cryptista were further lowered (Fig. 3B and F). In stark contrast, the exclusion of *P. bilix* enhanced the affinity between Rhodophyta and Cryptista (Fig. 3J). These results clearly indicated that, in the absence of Rhodelphidia and *M. maris*, *P. bilix* possesses a significant impact on the recovery of the monophyly of Archaeplastida by excluding Cryptista.

**Figure 3.**
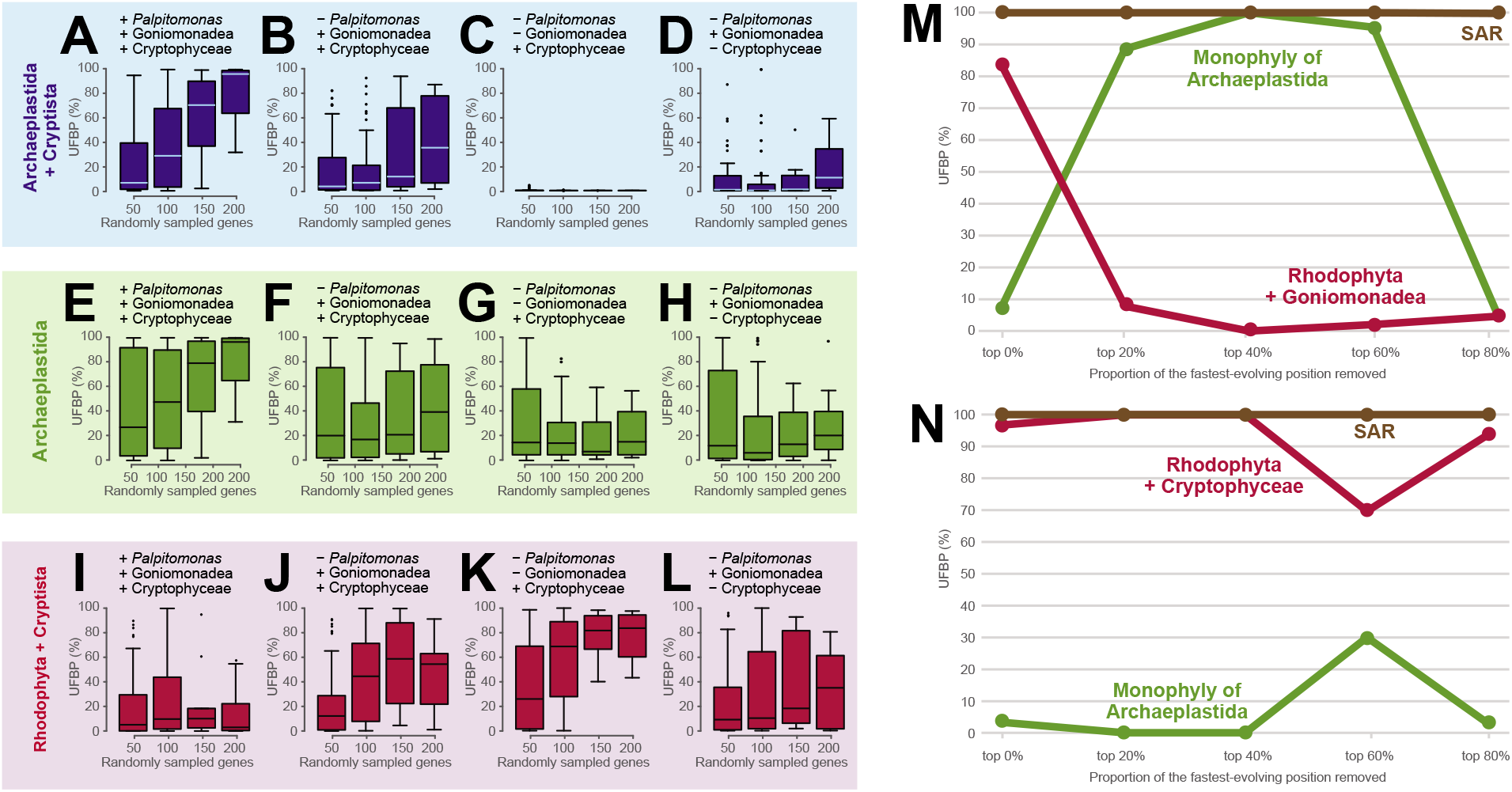
Analyses assessing the phylogenetic affinity of Rhodophyta to Cryptophyceae and/or Goniomonadea. **(A-L)** Analyses of the alignments generated by random gene sampling (RGS). We excluded both *Rhodelphis* spp. and *Microheliella maris* from the “rs50g,” “rs100g,” “rs150g,” and “rs200g” alignments, which were generated from the Diaph alignments (see Methods for the detail) and then subjected to the ultrafast bootstrap analyses using IQ-TREE 1.6.12. The ultrafast support values (UFBPs) for the sister relationship between Archaeplastida and Cryptista, the monophyly of Archaeplastida, and the union of Rhodophyta and Cryptista are presented as box-and-whisker plots (**A**), (**E**), and (**I**), respectively. The ultrafast bootstrap analyses on the rs50g, rs100g, rs150g, and rs200g alignments were repeated after further exclusion of *Palpitomonas bilix* (**B, F, and J**), *P. bilix* and Goniomonadea (**C, G, and K**), and *P. bilix* and Cryptophyceae (**D, H, and L**). The UFBPs shown in the plots described above are summarized in Table S4. **(M & N)** Analyses of the alignments processed by fast-evolving position removal (FPR). We modified the Diaph alignment in two ways, (i) the exclusion of *Rhodelphis* spp., *M. maris*, *P. bilix*, and Cryptophyceae and (ii) that of *Rhodelphis* spp., *M. maris*, *P. bilix*, and Goniomonadea. The two modified Diaph alignments were processed by FPR and further subjected to ultrafast bootstrap analyses. We plotted the UFBPs for the monophyly of the SAR clade (brown), those for the monophyly of Archaeplastida (green), and those for the grouping of Rhodophyta and Goniomonadea/Cryptophyceae (red).

We further analyzed the alignments in which Cryptista was represented solely by Cryptophyceae or Goniomonadea. The most drastic results were obtained from the analyses of the alignments in which the member of Cryptophyceae were the sole representatives of Cryptista (Figs. 3C, G, and K; see also Table S4 for the details). After the exclusion of the non-photosynthetic lineages in CAM clade (i.e., Rhodelphidia, *M. maris*, *P. bilix*, and Goniomonadea), the union of Cryptista (i.e., Cryptophyceae) and Rhodophyta appeared to dominate over the monophyly of Archaeplastida, particularly in the analyses of larger-size alignments (Figs. 3G and K). The analyses of rs200g alignments recovered the union of Cryptista and Rhodophyta with UFBPs ranging from 43.2 to 97.6% (Table S4). Of note, the suppression of UFBPs appeared to be much severer to the sister relationship between Archaeplastida and Cryptista than the monophyly of Archaeplastida—the UFBPs for the former grouping were 0 or nearly 0, regardless of the alignment size (Fig. 3C; Table S4). Comparing to the analyses considering Cryptophyceae as the sole representative of Cryptista, we observed only the mild suppression of the monophyly of Archaeplastida and that of the sister relationship between Archaeplastida and Cryptista in the analyses of the alignments in which Cryptista was represented by Goniomonadea (Figs. 3D and H; Table S4). These observations coincide with the affinity between Goniomonadea and Rhodophyta being less intense (Fig. 3L) than that between Cryptophyceae and Rhodophyta (Fig. 3K).

We revealed that Rhodophyta was attracted to Goniomonadea and Cryptophyceae erroneously in the ML phylogenies inferred from the alignments lacking Rhodelphidia, *M. maris*, and *P. bilix* (see above). In the global eukaryotic phylogeny, Rhodelphidia interrupts the branch leading to the Rhodophyta clade. Likewise, *M. maris* and *P. bilix*, both of which are basal branches of the Pancryptista clade, break the branch leading to the clade of Cryptophyceae and Goniomonadea (i.e., Cryptomonada). Thus, the grouping of Rhodophyta and Cryptomonada is most likely the phylogenetic artifact in which the two long branches, one exposed by the absence of Rhodelphidia and the other exposed by the absence of *M. maris* and *P. bilix*, attract to each other—long-branch attraction (LBA) artifact ^49^. Importantly, this phylogenetic artifact could not be overcome completely in the analyses of the alignments comprising at least 200 genes (Figs. 3J-L). If so, we anticipated that the putative phylogenetic artifact uniting Rhodophyta and Cryptomonada was enhanced further by the exclusion of Cryptophyceae (or Goniomonadea) (Figs. 3K and L), as this procedure extended the branch leading to the clade of Goniomonadea (or Cryptophyceae). The exclusion of Goniomonadea (i.e., Cryptophyceae were the sole representatives of Cryptomonada) appeared to enhance the putative phylogenetic artifact much greater degree than the exclusion of Cryptophyceae (i.e., Goniomonadea were the sole representatives of Cryptomonada) (compare Figs. 3C with D, Figs. 3G with H, and Figs. 3K and L). These results imply that the phylogenetic artifact uniting Rhodophyta and Cryptophyceae is substantially different from that uniting Rhodophyta and Goniomonadea. Altogether, we conclude that both size and taxon sampling, particularly the sampling of the members of CAM clade, in alignments heavily matter to reconstruct the monophyly of Archaeplastida and sister relationship between Archaeplastida and Pancryptista/Cryptista with confidence.

To pursue the reason why Rhodophyta is artifactually attracted to Cryptophyceae more severely than Goniomonadea (see above), we modified the Diaph alignment by excluding Rhodelphidia and all members of Pancryptista except Goniomonadea, and the resultant alignment was then subjected to FPR analysis (Fig. 3M). In the analysis of the alignment with full positions, the union of Goniomonadea and Rhodophyta received a UFBP of greater than 80%, while the monophyly of Archaeplastida was supported by a UFBP of smaller than 10%. The analyses of the alignments after removing the top 20-60% fastest-evolving positions drastically increased the UFBP for the monophyly of Archaeplastida (89-100%; green line in Fig. 4M), while the UFBP for the grouping of Rhodophyta and Goniomonadea was reduced to less than 10% (red line in Fig. 3M). Thus, we conclude that the union of Rhodophyta and Goniomonadea is the typical LBA artifact stemming from fast-evolving positions. We repeated the same analysis described above but substituted Goniomonadea with Cryptophyceae (Fig. 3N). Unexpectedly, the union of Rhodophyta and Cryptophyceae received UFBP of 98%, 100%, 100%, 70%, and 92% in the analyses after removal of top 20%, 40%, 60%, and 80% fastest-evolving positions, respectively (red line in Fig. 3N). The UFBP for the monophyly of Archaeplastida was less than 10%, except the analysis after removal of the top 60% fastest-evolving positions gave a UFBP of 30% (green line in Fig. 3N). These results strongly suggest that, in terms of dependency on fast-evolving positions, the phylogenetic artifact uniting Rhodophyta and Cryptophyceae is distinct from the typical LBA artifact uniting Rhodophyta and Goniomonadea.

### (d) The biological perspective on the signal uniting Rhodophyta and Cryptophyceae recovered in phylogenomic analyses

It is attractive to propose that the difference between the artifact uniting Rhodophyta and Cryptophyceae and that uniting Rhodophyta and Goniomonadea stems from the difference in lifestyle between the two closely related lineages in Cryptista. Goniomonadea is primarily heterotrophic and their nuclear genomes are free from endosymbiotic gene transfer (EGT) ^50^. Indeed, the series of the phylogenetic analyses described above demonstrated that the typical LBA was sufficient to explain the union of Rhodophyta and Goniomonadea (illustrated typically by Fig. 3M). In contrast, the extant member of Cryptophyceae possesses the plastids that were traced back to a red algal endosymbiont in the common ancestor of Cryptophyceae. During the red algal endosymbiont being transformed into a host-governed plastid, a number of genes had been transferred from the endosymbiont nucleus to the host nucleus. If a phylogenomic alignment contains genes acquired from the red algal endosymbiont, such genes are the source of the phylogenetic “signal” uniting Cryptophyceae and Rhodophyta. However, we selected the 319 genes, each of which showed no apparent sign of EGT in the corresponding single-gene phylogenetic analysis, for the phylogenomic analyses in this study. Additionally, we calculated the log-likelihoods (lnLs) of two identical tree topologies except for the position of Cryptophyceae—one bearing the monophyly of Archaeplastida (Tree 1) and the other bearing the grouping of Rhodophyta and Cryptophyceae (Tree 2)—for each of the 319 single-gene alignments (note that Rhodelphidia, *M. maris*, *P. bilix*, and Goniomonadea were omitted from the alignments) (Figs. S5A and S5B). The 319 single-gene alignments were sorted by the lnL difference between the two test trees (normalized by the alignment lengths) and the top 10 alignments, which prefer Tree 2 over Tree 1 most, were subjected individually to the standard ML phylogenetic analyses (Fig. S6, see also Table S5 for the details). Nevertheless, we detected any strong phylogenetic affinity between Rhodophyta and Cryptophyceae in none of the 10 ML single-gene analyses (Fig. S6, see also Table S5). These results cannot be explained by a simple scenario assuming that a subset of the ‘cryptophycean genes’ in the phylogenomic alignment was in fact acquired endosymbiotically from the red algal endosymbiont as briefly mentioned in Cavalier-Smith et al., (2015) ^30^. Rather, we here propose that a potentially large number of the cryptophycean nuclear genes (including those composed of the phylogenomic alignment) are the chimeras of the sequence inherited vertically beyond the red algal endosymbiosis and that acquired from the red algal endosymbiont. In each chimeric gene, the phylogenetic signal from the red alga-derived gene portion is likely insufficient to unite Rhodophyta and Cryptophyceae together in the single-gene analysis. However, when multiple chimeric genes in the cryptophycean nuclear genomes were included in a phylogenomic alignment, the phylogenetic signal from the red alga-derived gene portion becomes detectable as the union of Rhodophyta and Cryptista in the absence of Rhodelphidia and the basal branching taxa in Pancryptista, such as *M. maris* and *P. bilix*. Unfortunately, we additionally calculated the site-wise lnL differences between Trees 1 and 2, albeit no clear sign for the putative red algal gene fragments was detected (Fig. S7, see also Table S6).

## 4. Conclusion

In this work, we successfully deepen our understanding of the early evolution of eukaryotes. The phylogenomic analyses presented here demonstrated that *M. maris* is critical to understand the early evolution of Cryptista, as well as that of Archaeplastida. We also revealed that the deep branches of Archaeplastida and Cryptista—Rhodelphidia, *M. maris*, *P. bilix* (although not examined in this study, Picozoa most likely possesses the equivalent impact to the above-mentioned species/lineages, too)—are critical to suppress the cryptic and severe phylogenetic “signal” in cryptophycean genes.

## Data accessibility

The transcriptome data and genome data of strain *M. maris* were deposited in DDBJ Sequence Archive under the accession nos. DRAXXX and DRAYYY. The assembled transcriptomes and genome sequences of *M. maris* and phylogenetic alignments analyzed in this study are available from the Dryad Digital Repository.

## Acknowledgments

This study was supported by grants from the Japan Society for the Promotion of Science (18KK0203 & 19H03280) awarded to YI, (11J04684 & 17K19434) to AY, and was partially supported by the Tree of Life Project of University of Tsukuba. The phylogenetic analyses conducted in this work have been carried out under the ‘Interdisciplinary Computational Science Program’ in the Center for Computational Sciences, University of Tsukuba.

## Authors’ contributions

EY and AY participated in the design of the study, carried out the molecular lab work, carried out data analyses including phylogenetic analyses and drafted the manuscript; AI prepared the phylogenomic dataset; KK and TH assembled the RNA-seq data; YI participated in the design of the study and revised the manuscript. All authors gave final approval for publication and agree to be held accountable for the work performed therein.

## Declaration of interests

We declare no competing interests.

## Notes

### Competing Interest Statement

The authors have declared no competing interest.

