## Supplemental figures, S1-S7 for "Phylogenomics invokes the clade housing Cryptista, Archaeplastida, and *Microheliella maris*"

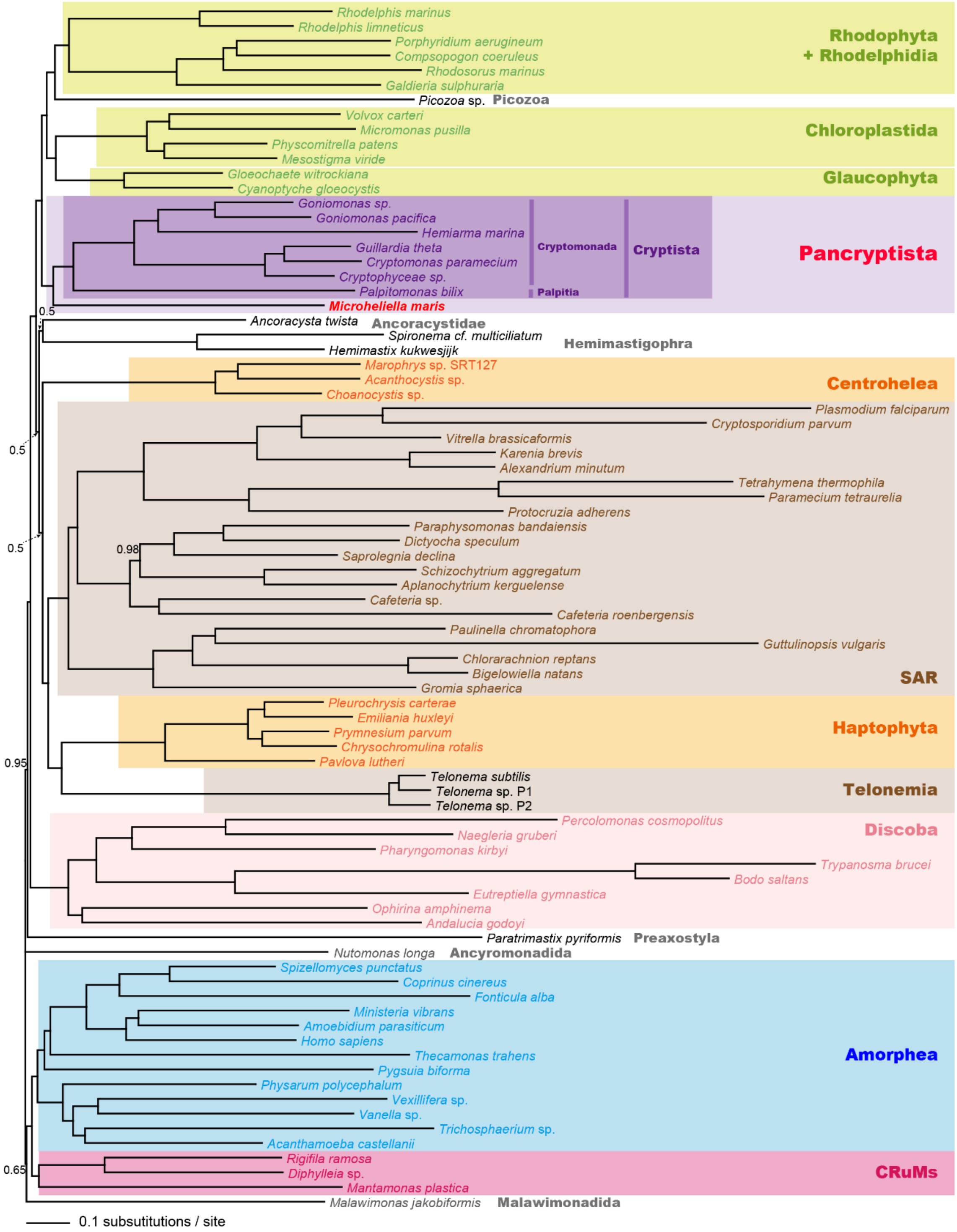


**Figure S1. Bayesian phylogeny inferred from the GlobE alignment.**

The tree topology and branch lengths were inferred from the GlobE alignment (319 genes; 88,592 amino acid positions in total) by Bayesian method. Bayesian posterior probabilities (BPPs) other than 1.0 are shown. Thus, all of the bipartitions with no value were supported by BPPs of 1.0.

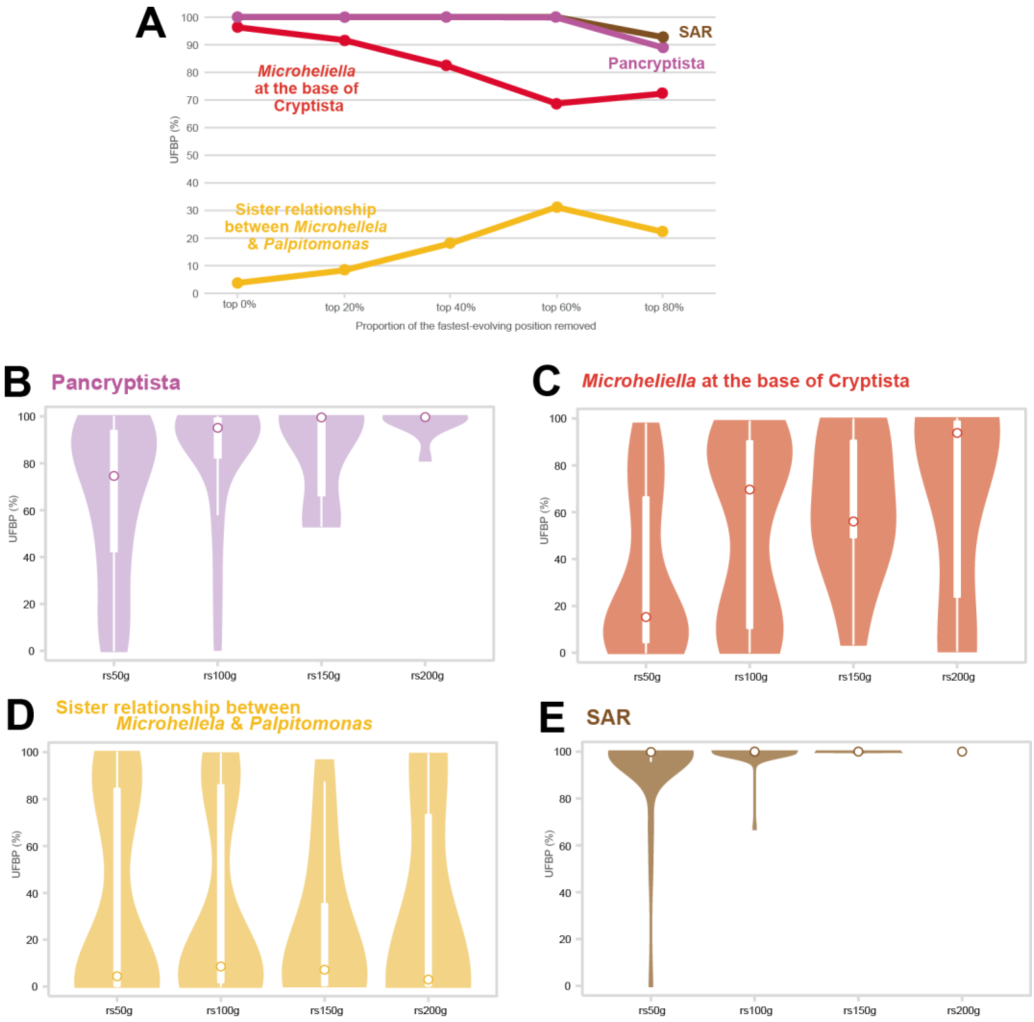


**Figure S2. Examinations of the phylogenetic position of *Microheliella maris* inferred from the GlobE alignment.**

**(A)** Analyses of GlobE alignment processed by fast-evolving position removal (FPR). We repeated ultra-fast bootstrap analyses using IQ-TREE 1.6.12 on the GlobE alignment after excluding no position, the top 20, 40, 60, and 80% fastest-evolving positions. The plots in brown, purple, red, and yellow indicate the ultra-fast bootstrap support values (UFBPs) for the monophyly of the SAR clade, the monophyly of Pancryptista, the basal position of *Microheliella maris* to the Cryptista clade, and the sister relationship between *M. maris* to *Palpitomonas bilix*, respectively. **(B)** Analyses of the alignment generated by random gene sampling (RGS). We sampled 50, 100, 150, and 200 genes randomly from the 319 genes in the GlobE alignment, concatenated into "rs50g," "rs100g," "rs150g," and "rs200g" alignments, and subjected to ultra-fast bootstrap analyses using IQ-TREE 1.6.12. We presented the UFBPs for the SAR clade, the monophyly of Pancryptista, the basal position of *M. maris* to the Cryptista, as kernel density estimate-box plots **(B)**, **(C), (D)**, and **(E)**, respectively. The UFBPs shown in the plots described above are summarized in Table S2.

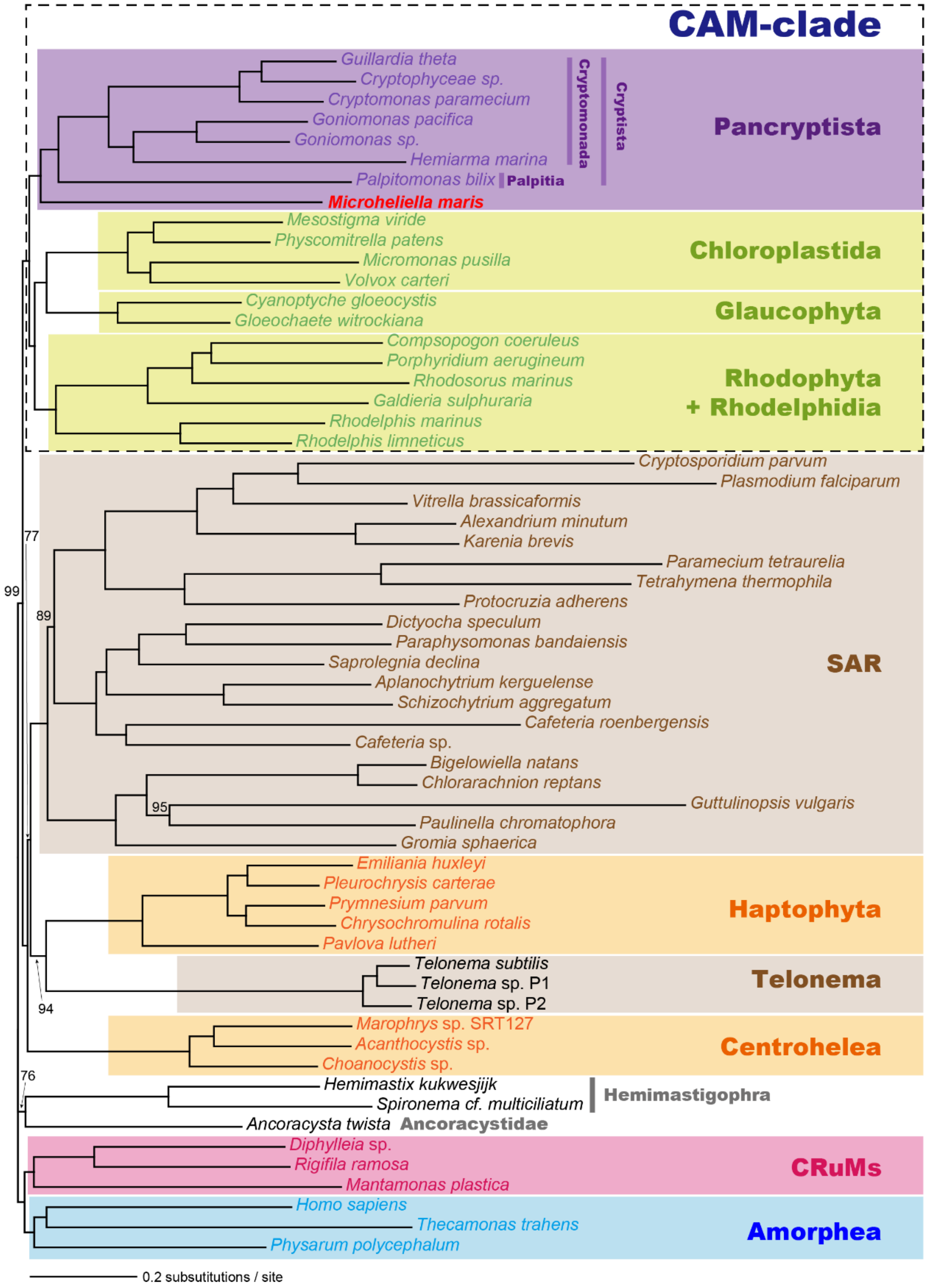


**Figure S3. Maximum-likelihood phylogeny inferred from the Diaph alignment.**

The tree topology and branch lengths were inferred from the Diaph alignment (319 genes; 88,592 amino acid positions in total) by the maximum-likelihood (ML) method. ML bootstrap support values (MLBPs) other than 100 are shown. Thus, all of the bipartitions with no value were supported by MLBPs of 100.

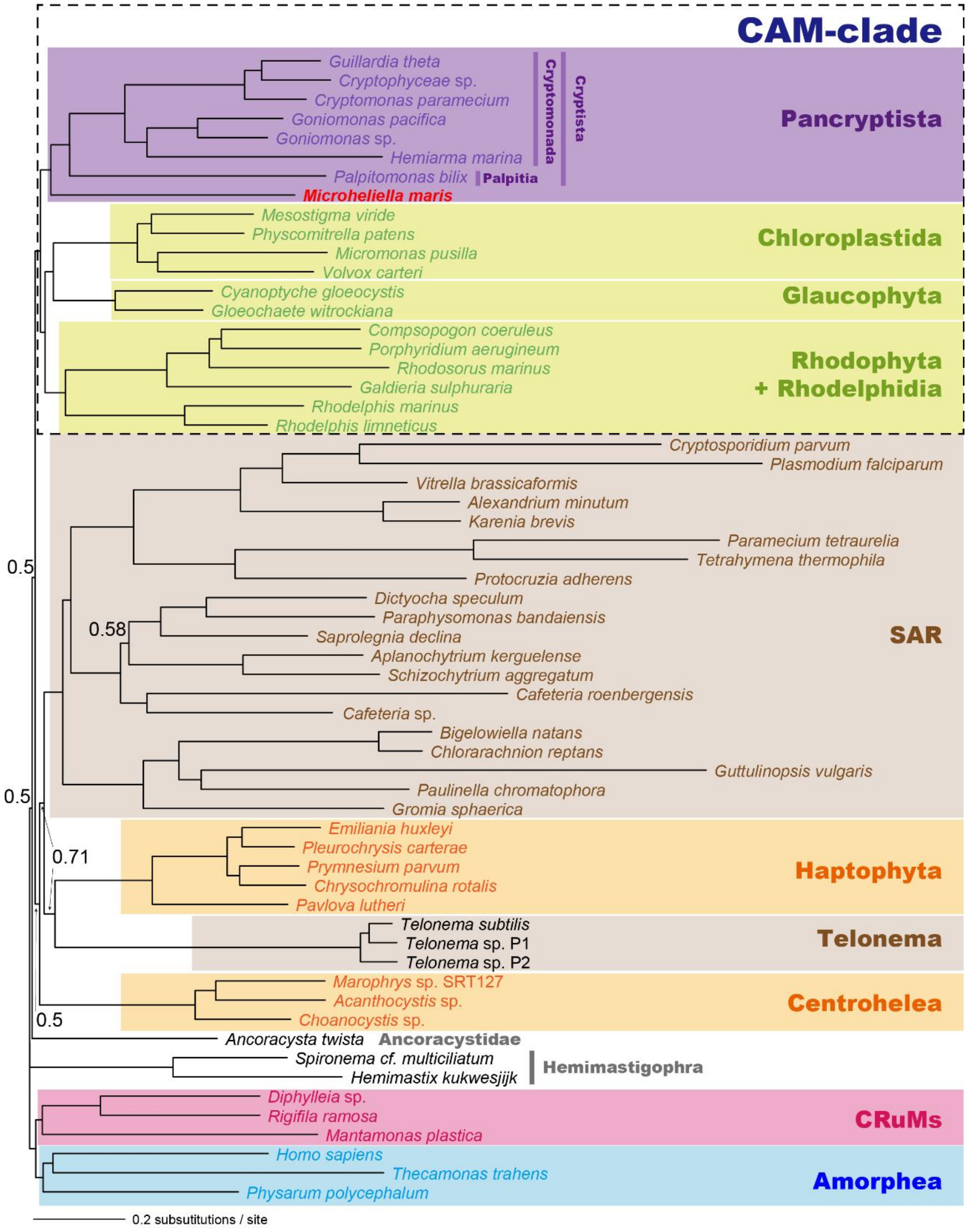


**Figure S4. Bayesian phylogeny inferred from the Diaph alignment.**

The tree topology and branch lengths were inferred from the Diaph alignment (319 genes; 88,592 amino acid positions in total) by Bayesian method. Bayesian posterior probabilities (BPPs) other than 1.0 are shown. Thus, all of the bipartitions with no value were supported by BPPs of 1.0.

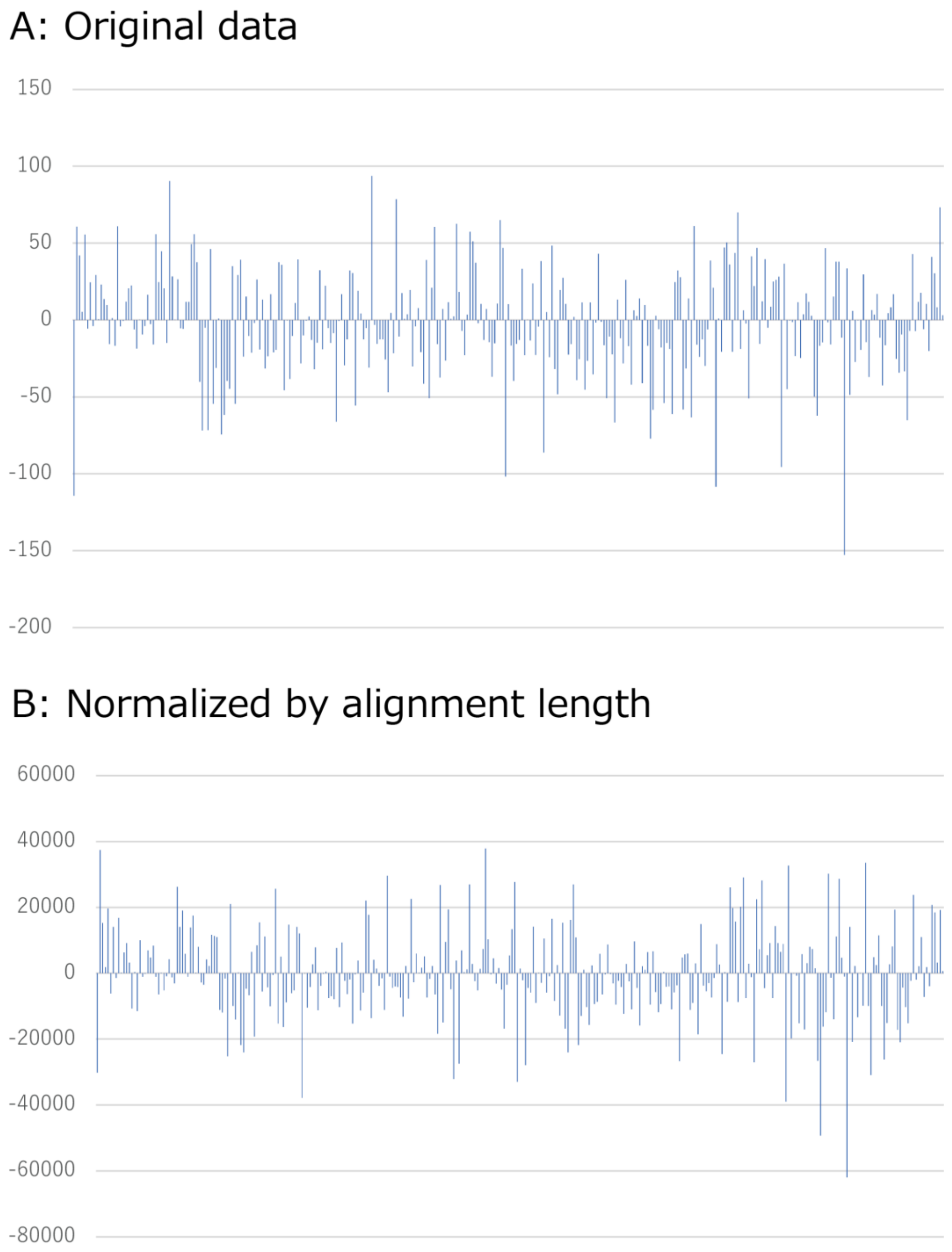


**Figure S5. Gene-wise comparison between two competing hypotheses for the position of Cryptophyceae**.
We prepared two trees in which only the position of Cryptophyceae is different. "Tree 1" is identical to the maximum-likelihood tree inferred from Daiph alignment (Fig. S3), except *Microheliella maris*, *Palpitomonas bilix*, Goniomonadea, and Rhodelphidia were pruned. We modified Tree 1 by forcing Cryptophyceae to branch with Rhodophyta directly to prepare "Tree 2." For each of the 319 single-gene alignments, the log-likelihoods (lnLs) of the two test trees were calculated with the LG + Γ + F model using IQTREE 1.6.12. (A) The gene-wise lnL differences obtained by subtracting the lnL of Tree 2 from that of Tree 1. (B) The gene-wise lnL differences normalized by alignment size. The lnL differences are sorted from left to right on the horizontal axis according to the gene numbers given in Table S5.

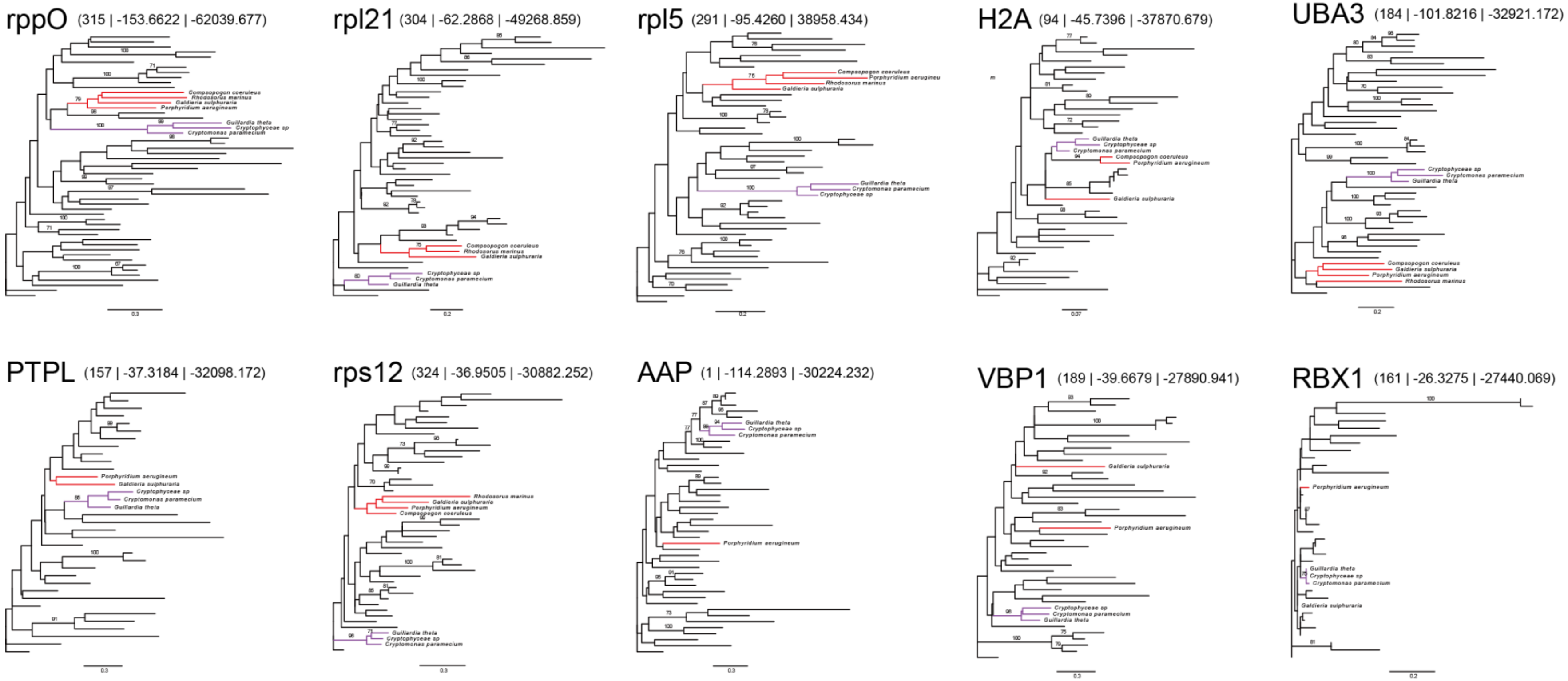


**Figure S6. Maximum-likelihood phylogenies inferred from the single-gene alignments that showed the top 10 largest gene-wise log-likelihood differences between Trees 1 and 2.**
Based on the normalized gene-wise log-likelihood (lnL) differences between Trees 1 and 2 (see Fig. S5B), we identified the top 10 genes that preferred Tree 2 over Tree 1 and subjected them individually to the maximum likelihood (ML) analyses with LG + Γ + F model using IQ-TREE 1.6.12. The statistical support values for bipartitions are calculated by the UFBOOT approximation in IQ-TREE (1,000 replicates). Only ultra-fast bootstrap support values are shown. For each of the 10 single-gene phylogenies, the gene number given in Tables S1 and S6, gene-wise lnL difference, and normalized gene-wise lnL difference are given in parentheses (the three figures are separated by dividers). The branches in red and those in purple indicate Rhodophyta and Cryptophyceae, respectively.

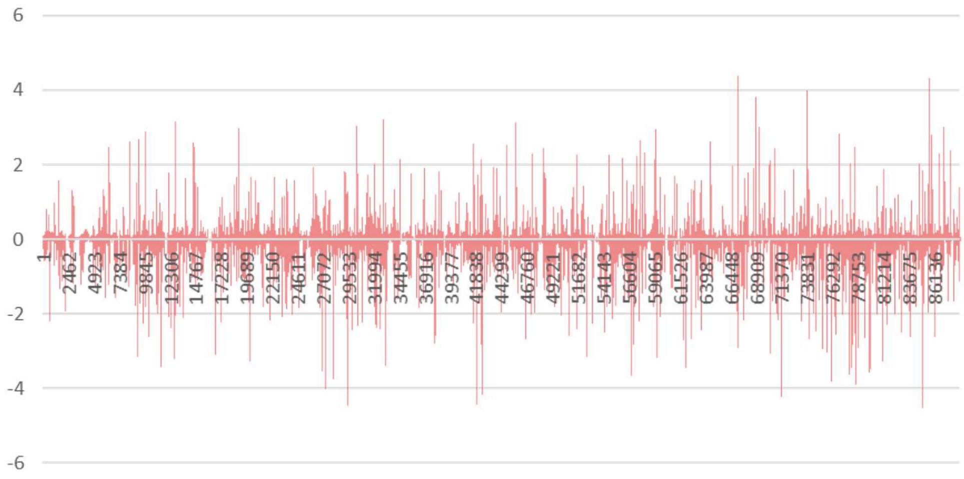


**Figure S7. Site-wise comparison of log-likelihood between two competing hypotheses for the position of Cryptophyceae**.

We calculated the log-likelihood of each alignment positions (site-lnL) in the 319-gene alignment over Trees 1 and 2 (see the legend of Figure S5) with the LG + Γ + F model using IQTREE 1.6.12. For each alignment position, we subtracted the site-lnL calculated over Tree 2 from the corresponding value calculated over Tree 1. The site-lnL differences are sorted from left to right on the horizontal axis according to the site numbers given in Table S6.
