## Supplemental Table 2 for "Phylogenomics invokes the clade housing Cryptista, Archaeplastida, and *Microheliella maris*"

| | rs50g | rs50g | rs50g | rs50g | rs100g | rs100g | rs100g | rs100g | rs150g | rs150g | rs150g | rs150g | rs200g | rs200g | rs200g | rs200g |
|---|---|---|---|---|---|---|---|---|---|---|---|---|---|---|---|---|
| Replicate # | SAR | Pancryptista | Microheliella at the base of Cryptista | Sister relationship between Microheliella and Palpitomonas | SAR | Pancryptista | Microheliella at the base of Cryptista | Sister relationship between Microheliella and Palpitomonas | SAR | Pancryptista | Microheliella at the base of Cryptista | Sister relationship between Microheliella and Palpitomonas | SAR | Pancryptista | Microheliella at the base of Cryptista | Sister relationship between Microheliella and Palpitomonas |
| 0 | 999 | 979 | 977 | 2 | 1000 | 820 | 804 | 16 | 1000 | 999 | 996 | 3 | 1000 | 993 | 48 | 950 |
| 1 | 1000 | 917 | 894 | 23 | 1000 | 989 | 270 | 727 | 1000 | 567 | 561 | 6 | 1000 | 1000 | 990 | 10 |
| 2 | 926 | 732 | 675 | 57 | 998 | 814 | 724 | 90 | 1000 | 997 | 36 | 964 | 1000 | 1000 | 1000 | 0 |
| 3 | 1000 | 883 | 7 | 991 | 1000 | 541 | 530 | 14 | 1000 | 1000 | 896 | 104 | 1000 | 939 | 8 | 992 |
| 4 | 994 | 1000 | 44 | 956 | 1000 | 809 | 771 | 38 | 999 | 995 | 909 | 86 | 1000 | 992 | 992 | 0 |
| 5 | 979 | 927 | 49 | 932 | 1000 | 1000 | 39 | 961 | 1000 | 565 | 560 | 5 | 1000 | 998 | 956 | 42 |
| 6 | 992 | 898 | 195 | 709 | 1000 | 40 | 40 | 0 | 1000 | 958 | 545 | 435 | 1000 | 1000 | 921 | 79 |
| 7 | 1000 | 129 | 128 | 1 | 1000 | 648 | 638 | 25 | 1000 | 1000 | 998 | 2 | 1000 | 1000 | 48 | 952 |
| 8 | 959 | 974 | 3 | 997 | 1000 | 986 | 18 | 968 | 1000 | 532 | 475 | 57 | 1000 | 996 | 979 | 18 |
| 9 | 1000 | 886 | 885 | 1 | 1000 | 954 | 945 | 14 | 998 | 1000 | 364 | 636 | 1000 | 813 | 811 | 2 |
| 10 | 991 | 952 | 0 | 1000 | 1000 | 980 | 886 | 94 | | | | | | | | |
| 11 | 1000 | 328 | 277 | 57 | 1000 | 5 | 5 | 0 | | | | | | | | |
| 12 | 991 | 444 | 22 | 971 | 1000 | 998 | 97 | 901 | | | | | | | | |
| 13 | 1000 | 760 | 740 | 20 | 1000 | 999 | 14 | 985 | | | | | | | | |
| 14 | 1000 | 48 | 47 | 1 | 1000 | 972 | 966 | 6 | | | | | | | | |
| 15 | 1000 | 180 | 104 | 76 | 999 | 795 | 746 | 49 | | | | | | | | |
| 16 | 1000 | 993 | 29 | 968 | 1000 | 990 | 988 | 2 | | | | | | | | |
| 17 | 998 | 627 | 620 | 7 | 1000 | 881 | 689 | 192 | | | | | | | | |
| 18 | 1000 | 987 | 208 | 791 | 1000 | 943 | 807 | 165 | | | | | | | | |
| 19 | 1000 | 578 | 7 | 897 | 999 | 271 | 254 | 17 | | | | | | | | |
| 20 | 993 | 859 | 849 | 10 | 998 | 1000 | 83 | 917 | | | | | | | | |
| 21 | 83 | 844 | 82 | 786 | 1000 | 561 | 477 | 84 | | | | | | | | |
| 22 | 979 | 381 | 157 | 234 | 1000 | 992 | 892 | 100 | | | | | | | | |
| 23 | 977 | 979 | 2 | 979 | 1000 | 972 | 945 | 30 | | | | | | | | |
| 24 | 1000 | 440 | 440 | 0 | 1000 | 997 | 950 | 47 | | | | | | | | |
| 25 | 1000 | 422 | 422 | 0 | 698 | 993 | 2 | 994 | | | | | | | | |
| 26 | 1000 | 835 | 834 | 1 | 1000 | 946 | 945 | 1 | | | | | | | | |
| 27 | 991 | 986 | 35 | 953 | 1000 | 220 | 198 | 22 | | | | | | | | |
| 28 | 999 | 965 | 963 | 2 | 1000 | 560 | 287 | 576 | | | | | | | | |
| 29 | 976 | 963 | 35 | 962 | 1000 | 881 | 843 | 38 | | | | | | | | |
| 30 | 987 | 984 | 127 | 862 | 1000 | 981 | 121 | 862 | | | | | | | | |
| 31 | 948 | 702 | 188 | 517 | 990 | 945 | 924 | 21 | | | | | | | | |
| 32 | 667 | 946 | 944 | 2 | 1000 | 949 | 948 | 1 | | | | | | | | |
| 33 | 994 | 888 | 143 | 746 | 1000 | 844 | 841 | 3 | | | | | | | | |
| 34 | 987 | 909 | 878 | 31 | 1000 | 972 | 967 | 5 | | | | | | | | |
| 35 | 532 | 868 | 51 | 911 | 989 | 959 | 96 | 867 | | | | | | | | |
| 36 | 1000 | 3 | 0 | 5 | 1000 | 885 | 736 | 149 | | | | | | | | |
| 37 | 742 | 670 | 209 | 536 | 1000 | 365 | 321 | 44 | | | | | | | | |
| 38 | 1000 | 941 | 189 | 763 | 1000 | 994 | 121 | 875 | | | | | | | | |
| 39 | 1000 | 494 | 486 | 8 | 1000 | 927 | 925 | 2 | | | | | | | | |
| 40 | 998 | 1 | 1 | 0 | 1000 | 973 | 125 | 852 | | | | | | | | |
| 41 | 1000 | 122 | 118 | 4 | 999 | 926 | 43 | 931 | | | | | | | | |
| 42 | 984 | 139 | 139 | 0 | 1000 | 1000 | 942 | 58 | | | | | | | | |
| 43 | 999 | 386 | 148 | 240 | 1000 | 972 | 801 | 171 | | | | | | | | |
| 44 | 997 | 641 | 636 | 5 | 1000 | 970 | 704 | 266 | | | | | | | | |
| 45 | 999 | 91 | 91 | 0 | 1000 | 1000 | 9 | 991 | | | | | | | | |
| 46 | 1000 | 715 | 708 | 9 | 1000 | 946 | 39 | 961 | | | | | | | | |
| 47 | 1000 | 720 | 718 | 2 | 1000 | 933 | 907 | 26 | | | | | | | | |
| 48 | 1000 | 0 | 0 | 0 | 1000 | 1000 | 913 | 87 | | | | | | | | |
| 49 | 971 | 690 | 673 | 17 | 1000 | 999 | 100 | 899 | | | | | | | | |
