## Supplemental Table 3 for "Phylogenomics invokes the clade housing Cryptista, Archaeplastida, and *Microheliella maris*"

| | rs50g | rs50g | rs50g | rs100g | rs100g | rs100g |
|---|---|---|---|---|---|---|
| Replicate # | Sister relationship between Archaeplastida and Pancryptista | Archaeplastida | Pancryptista | Sister relationship between Archaeplastida and Pancryptista | Archaeplastida | Pancryptista |
| 0 | 593 | 607 | 1000 | 277 | 982 | 791 |
| 1 | 602 | 778 | 749 | 972 | 983 | 999 |
| 2 | 545 | 797 | 815 | 768 | 996 | 950 |
| 3 | 811 | 993 | 999 | 872 | 914 | 995 |
| 4 | 926 | 944 | 1000 | 510 | 519 | 963 |
| 5 | 168 | 910 | 658 | 934 | 934 | 1000 |
| 6 | 832 | 843 | 994 | 205 | 863 | 256 |
| 7 | 159 | 161 | 200 | 898 | 957 | 957 |
| 8 | 736 | 896 | 1000 | 991 | 993 | 1000 |
| 9 | 7 | 7 | 911 | 917 | 973 | 975 |
| 10 | 899 | 1000 | 1000 | 894 | 997 | 928 |
| 11 | 100 | 922 | 341 | 655 | 712 | 986 |
| 12 | 554 | 674 | 995 | 818 | 930 | 998 |
| 13 | 148 | 186 | 889 | 522 | 587 | 1000 |
| 14 | 35 | 800 | 426 | 485 | 726 | 976 |
| 15 | 103 | 856 | 437 | 539 | 961 | 572 |
| 16 | 683 | 753 | 998 | 718 | 877 | 994 |
| 17 | 278 | 754 | 346 | 282 | 645 | 905 |
| 18 | 20 | 25 | 996 | 901 | 1000 | 906 |
| 19 | 837 | 996 | 963 | 169 | 185 | 674 |
| 20 | 601 | 683 | 991 | 36 | 763 | 991 |
| 21 | 653 | 718 | 958 | 633 | 767 | 998 |
| 22 | 4 | 463 | 347 | 753 | 901 | 999 |
| 23 | 58 | 515 | 966 | 857 | 857 | 994 |
| 24 | 9 | 9 | 346 | 601 | 773 | 766 |
| 25 | 581 | 925 | 636 | 278 | 913 | 980 |
| 26 | 6 | 8 | 330 | 544 | 983 | 973 |
| 27 | 795 | 855 | 864 | 305 | 872 | 662 |
| 28 | 118 | 971 | 694 | 922 | 947 | 991 |
| 29 | 107 | 447 | 941 | 480 | 577 | 974 |
| 30 | 690 | 763 | 985 | 280 | 280 | 997 |
| 31 | 826 | 859 | 957 | 962 | 974 | 990 |
| 32 | 754 | 768 | 973 | 358 | 996 | 979 |
| 33 | 119 | 826 | 913 | 738 | 773 | 919 |
| 34 | 496 | 945 | 852 | 668 | 817 | 967 |
| 35 | 718 | 940 | 950 | 970 | 974 | 999 |
| 36 | 83 | 624 | 123 | 813 | 838 | 994 |
| 37 | 286 | 952 | 990 | 520 | 537 | 534 |
| 38 | 199 | 1000 | 486 | 179 | 583 | 993 |
| 39 | 32 | 176 | 317 | 949 | 1000 | 950 |
| 40 | 580 | 804 | 914 | 89 | 91 | 904 |
| 41 | 194 | 229 | 697 | 939 | 969 | 979 |
| 42 | 20 | 907 | 326 | 850 | 851 | 999 |
| 43 | 218 | 940 | 310 | 561 | 602 | 889 |
| 44 | 18 | 18 | 678 | 28 | 56 | 917 |
| 45 | 0 | 824 | 53 | 500 | 980 | 1000 |
| 46 | 654 | 780 | 739 | 837 | 978 | 982 |
| 47 | 432 | 873 | 960 | 781 | 796 | 965 |
| 48 | 44 | 121 | 61 | 779 | 808 | 999 |
| 49 | 688 | 739 | 958 | 532 | 999 | 998 |

| | rs150g | rs150g | rs150g | rs200g | rs200g | rs200g |
|---|---|---|---|---|---|---|
| Replicate # | Sister relationship between Archaeplastida and Pancryptista | Archaeplastida | Pancryptista | Sister relationship between Archaeplastida and Pancryptista | Archaeplastida | Pancryptista |
| 0 | 911 | 983 | 1000 | 996 | 1000 | 1000 |
| 1 | 914 | 1000 | 949 | 977 | 993 | 994 |
| 2 | 982 | 1000 | 1000 | 993 | 1000 | 998 |
| 3 | 627 | 1000 | 767 | 947 | 1000 | 999 |
| 4 | 887 | 889 | 1000 | 994 | 1000 | 1000 |
| 5 | 973 | 979 | 1000 | 992 | 999 | 1000 |
| 6 | 859 | 869 | 1000 | 989 | 991 | 998 |
| 7 | 997 | 998 | 1000 | 985 | 989 | 1000 |
| 8 | 915 | 996 | 948 | 955 | 998 | 993 |
| 9 | 928 | 960 | 1000 | 973 | 1000 | 995 |

| Classification | | | Taxon sampling | |
|---|---|---|---|---|
| Archaeplastida | | | *Rhodelphis* spp. | ✓ |
| Pancryptista | | | *Microheliella maris* | ✓ |
| Pancryptista | Cryptista | | *Palpitomonas bilix* | ✓ |
| Pancryptista | Cryptista | Cryptomonada | Cryptophyceae | ✓ |
| Pancryptista | Cryptista | Cryptomonada | Goniomonadea | ✓ |

| Replicate # | rs50g Sister relationship between Archaeplastida and Pancryptista | rs50g Archaeplastida | rs50g Pancryptista | rs100g Sister relationship between Archaeplastida and Pancryptista | rs100g Archaeplastida | rs100g Pancryptista |
|---|---|---|---|---|---|---|
| 0 | 397 | 467 | 999 | 112 | 943 | 755 |
| 1 | 290 | 386 | 572 | 976 | 988 | 1000 |
| 2 | 63 | 74 | 521 | 443 | 914 | 862 |
| 3 | 766 | 912 | 992 | 849 | 876 | 929 |
| 4 | 903 | 917 | 998 | 0 | 0 | 697 |
| 5 | 137 | 837 | 814 | 941 | 941 | 979 |
| 6 | 945 | 968 | 994 | 114 | 482 | 194 |
| 7 | 418 | 1000 | 531 | 661 | 691 | 851 |
| 8 | 371 | 986 | 994 | 944 | 955 | 969 |
| 9 | 5 | 5 | 91 | 694 | 784 | 812 |
| 10 | 225 | 916 | 992 | 950 | 986 | 999 |
| 11 | 79 | 980 | 252 | 436 | 504 | 910 |
| 12 | 599 | 999 | 995 | 672 | 707 | 997 |
| 13 | 1 | 25 | 135 | 463 | 569 | 946 |
| 14 | 13 | 855 | 301 | 25 | 148 | 964 |
| 15 | 12 | 471 | 289 | 289 | 517 | 451 |
| 16 | 412 | 706 | 990 | 334 | 416 | 698 |
| 17 | 0 | 9 | 542 | 24 | 85 | 791 |
| 18 | 9 | 18 | 965 | 799 | 976 | 832 |
| 19 | 350 | 977 | 876 | 24 | 28 | 358 |
| 20 | 4 | 17 | 994 | 15 | 416 | 788 |
| 21 | 288 | 453 | 711 | 801 | 867 | 943 |
| 22 | 0 | 18 | 309 | 632 | 998 | 957 |
| 23 | 24 | 249 | 962 | 236 | 236 | 279 |
| 24 | 0 | 0 | 177 | 537 | 857 | 700 |
| 25 | 227 | 258 | 486 | 361 | 882 | 997 |
| 26 | 0 | 14 | 353 | 605 | 762 | 936 |
| 27 | 802 | 985 | 987 | 140 | 231 | 810 |
| 28 | 43 | 969 | 706 | 632 | 697 | 947 |
| 29 | 626 | 845 | 995 | 19 | 25 | 370 |
| 30 | 58 | 70 | 480 | 111 | 126 | 798 |
| 31 | 788 | 857 | 945 | 966 | 977 | 987 |
| 32 | 563 | 575 | 998 | 365 | 612 | 994 |
| 33 | 40 | 905 | 796 | 423 | 460 | 630 |
| 34 | 505 | 996 | 862 | 445 | 560 | 812 |
| 35 | 558 | 675 | 892 | 519 | 526 | 613 |
| 36 | 0 | 21 | 146 | 301 | 306 | 456 |
| 37 | 434 | 565 | 881 | 991 | 996 | 996 |
| 38 | 31 | 998 | 423 | 536 | 785 | 917 |
| 39 | 23 | 64 | 411 | 907 | 1000 | 908 |
| 40 | 403 | 463 | 726 | 145 | 162 | 552 |
| 41 | 117 | 220 | 177 | 884 | 1000 | 912 |
| 42 | 8 | 919 | 160 | 834 | 835 | 845 |
| 43 | 112 | 990 | 309 | 36 | 86 | 618 |
| 44 | 0 | 0 | 851 | 9 | 40 | 295 |
| 45 | 0 | 672 | 51 | 377 | 992 | 1000 |
| 46 | 161 | 389 | 344 | 690 | 851 | 918 |
| 47 | 51 | 923 | 939 | 66 | 66 | 255 |
| 48 | 0 | 26 | 30 | 583 | 802 | 982 |
| 49 | 632 | 769 | 839 | 777 | 998 | 999 |

| Replicate # | rs150g Sister relationship between Archaeplastida and Pancryptista | rs150g Archaeplastida | rs150g Pancryptista | rs200g Sister relationship between Archaeplastida and Pancryptista | rs200g Archaeplastida | rs200g Pancryptista |
|---|---|---|---|---|---|---|
| 0 | 652 | 733 | 796 | 998 | 1000 | 1000 |
| 1 | 607 | 644 | 612 | 918 | 939 | 939 |
| 2 | 973 | 1000 | 1000 | 985 | 1000 | 1000 |
| 3 | 588 | 998 | 735 | 945 | 1000 | 999 |
| 4 | 819 | 822 | 941 | 898 | 901 | 972 |
| 5 | 902 | 911 | 912 | 980 | 1000 | 999 |
| 6 | 953 | 979 | 993 | 982 | 995 | 1000 |
| 7 | 905 | 906 | 917 | 959 | 989 | 1000 |
| 8 | 897 | 924 | 982 | 897 | 944 | 964 |
| 9 | 826 | 992 | 993 | 873 | 916 | 899 |

| Classification | | | Taxon sampling | |
|---|---|---|---|---|
| Archaeplastida | | | *Rhodelphis* spp. | |
| Pancryptista | | | *Microheliella maris* | ✓ |
| Pancryptista | Cryptista | | *Palpitomonas bilix* | ✓ |
| Pancryptista | Cryptista | Cryptomonada | Cryptophyceae | ✓ |
| Pancryptista | Cryptista | Cryptomonada | Goniomonadea | ✓ |

| | rs50g | rs50g | rs50g | rs100g | rs100g | rs100g |
|---|---|---|---|---|---|---|
| Replicate # | Sister relationship between Archaeplastida and Cryptista | Archaeplastida | Cryptista | Sister relationship between Archaeplastida and Cryptista | Archaeplastida | Cryptista |
| 0 | 369 | 390 | 1000 | 268 | 915 | 1000 |
| 1 | 126 | 128 | 1000 | 770 | 927 | 1000 |
| 2 | 237 | 258 | 1000 | 800 | 996 | 1000 |
| 3 | 859 | 968 | 993 | 600 | 700 | 1000 |
| 4 | 807 | 828 | 1000 | 169 | 185 | 1000 |
| 5 | 154 | 957 | 994 | 665 | 669 | 1000 |
| 6 | 309 | 310 | 1000 | 361 | 731 | 1000 |
| 7 | 743 | 942 | 1000 | 759 | 769 | 1000 |
| 8 | 677 | 807 | 969 | 938 | 938 | 1000 |
| 9 | 2 | 2 | 1000 | 541 | 970 | 1000 |
| 10 | 493 | 585 | 999 | 937 | 996 | 1000 |
| 11 | 50 | 864 | 215 | 468 | 532 | 1000 |
| 12 | 281 | 292 | 999 | 625 | 824 | 1000 |
| 13 | 98 | 657 | 999 | 317 | 477 | 1000 |
| 14 | 430 | 570 | 1000 | 442 | 500 | 1000 |
| 15 | 262 | 596 | 1000 | 288 | 315 | 1000 |
| 16 | 572 | 623 | 1000 | 487 | 693 | 1000 |
| 17 | 731 | 753 | 1000 | 216 | 361 | 1000 |
| 18 | 5 | 8 | 1000 | 958 | 981 | 1000 |
| 19 | 325 | 1000 | 993 | 0 | 0 | 1000 |
| 20 | 224 | 246 | 1000 | 11 | 648 | 1000 |
| 21 | 129 | 129 | 1000 | 632 | 805 | 1000 |
| 22 | 4 | 255 | 999 | 691 | 773 | 1000 |
| 23 | 16 | 674 | 997 | 256 | 256 | 1000 |
| 24 | 49 | 98 | 1000 | 180 | 182 | 1000 |
| 25 | 537 | 656 | 1000 | 398 | 734 | 1000 |
| 26 | 29 | 61 | 1000 | 273 | 938 | 1000 |
| 27 | 427 | 990 | 995 | 472 | 612 | 1000 |
| 28 | 85 | 980 | 1000 | 454 | 468 | 1000 |
| 29 | 247 | 381 | 998 | 221 | 497 | 1000 |
| 30 | 532 | 565 | 1000 | 92 | 92 | 1000 |
| 31 | 704 | 720 | 1000 | 935 | 937 | 1000 |
| 32 | 660 | 711 | 1000 | 548 | 993 | 1000 |
| 33 | 26 | 937 | 1000 | 220 | 220 | 1000 |
| 34 | 681 | 737 | 1000 | 470 | 586 | 1000 |
| 35 | 654 | 891 | 996 | 961 | 977 | 1000 |
| 36 | 667 | 706 | 1000 | 576 | 636 | 1000 |
| 37 | 376 | 840 | 1000 | 771 | 771 | 1000 |
| 38 | 131 | 998 | 998 | 222 | 643 | 1000 |
| 39 | 101 | 122 | 1000 | 904 | 1000 | 1000 |
| 40 | 215 | 233 | 1000 | 36 | 37 | 1000 |
| 41 | 21 | 21 | 1000 | 943 | 955 | 1000 |
| 42 | 363 | 845 | 1000 | 675 | 675 | 1000 |
| 43 | 536 | 900 | 1000 | 117 | 117 | 1000 |
| 44 | 0 | 0 | 1000 | 23 | 82 | 1000 |
| 45 | 2 | 689 | 1000 | 433 | 913 | 1000 |
| 46 | 591 | 666 | 1000 | 850 | 931 | 1000 |
| 47 | 657 | 936 | 1000 | 82 | 85 | 1000 |
| 48 | 52 | 54 | 1000 | 427 | 464 | 1000 |
| 49 | 134 | 138 | 1000 | 297 | 999 | 1000 |

| | rs150g | rs150g | rs150g | rs200g | rs200g | rs200g |
|---|---|---|---|---|---|---|
| Replicate # | Sister relationship between Archaeplastida and Cryptista | Archaeplastida | Cryptista | Sister relationship between Archaeplastida and Cryptista | Archaeplastida | Cryptista |
| 0 | 685 | 967 | 1000 | 995 | 998 | 1000 |
| 1 | 720 | 738 | 1000 | 743 | 755 | 1000 |
| 2 | 963 | 996 | 1000 | 972 | 997 | 1000 |
| 3 | 716 | 961 | 1000 | 915 | 926 | 1000 |
| 4 | 279 | 1000 | 1000 | 906 | 928 | 1000 |
| 5 | 786 | 801 | 1000 | 976 | 992 | 1000 |
| 6 | 241 | 1000 | 1000 | 939 | 939 | 1000 |
| 7 | 936 | 940 | 1000 | 929 | 932 | 1000 |
| 8 | 711 | 858 | 1000 | 889 | 925 | 1000 |
| 9 | 783 | 829 | 1000 | 951 | 992 | 1000 |

<table>
<tr><th colspan="3">Classification</th><th colspan="2">Taxon sampling</th></tr>
<tr><td colspan="3">Archaeplastida</td><td>Rhodelphis spp.</td><td>✓</td></tr>
<tr><td rowspan="4">Pancryptista</td><td colspan="2"></td><td>Microheliella maris</td><td></td></tr>
<tr><td rowspan="3">Cryptista</td><td></td><td>Palpitomonas biliix</td><td>✓</td></tr>
<tr><td rowspan="2">Cryptomonada</td><td>Cryptophyceae</td><td>✓</td></tr>
<tr><td>Goniomonadea</td><td>✓</td></tr>
</table>
