## Supplemental Table 4 for "Phylogenomics invokes the clade housing Cryptista, Archaeplastida, and *Microheliella maris*"

| | rs50g | rs50g | rs50g | rs100g | rs100g | rs100g | rs150g | rs150g | rs150g | rs200g | rs200g | rs200g |
|---|---|---|---|---|---|---|---|---|---|---|---|---|
| Replicate # | Sister relationship between Archaeplastida and Cryptista | Archaeplastida | Rhodophyta + Cryptista | Sister relationship between Archaeplastida and Cryptista | Archaeplastida | Rhodophyta + Cryptista | Sister relationship between Archaeplastida and Cryptista | Archaeplastida | Rhodophyta + Cryptista | Sister relationship between Archaeplastida and Cryptista | Archaeplastida | Rhodophyta + Cryptista |
| 0 | 129 | 167 | 8 | 214 | 873 | 0 | 623 | 814 | 167 | 977 | 983 | 17 |
| 1 | 42 | 45 | 4 | 867 | 941 | 56 | 281 | 287 | 182 | 630 | 651 | 220 |
| 2 | 40 | 43 | 0 | 543 | 778 | 541 | 981 | 998 | 2 | 976 | 1000 | 0 |
| 3 | 656 | 784 | 15 | 494 | 541 | 14 | 833 | 1000 | 0 | 983 | 996 | 4 |
| 4 | 753 | 793 | 855 | 0 | 0 | 434 | 19 | 21 | 936 | 312 | 314 | 567 |
| 5 | 529 | 821 | 194 | 895 | 900 | 26 | 935 | 949 | 45 | 985 | 1000 | 0 |
| 6 | 938 | 940 | 1 | 156 | 241 | 48 | 363 | 399 | 599 | 950 | 953 | 42 |
| 7 | 586 | 999 | 833 | 159 | 159 | 384 | 770 | 772 | 56 | 842 | 855 | 144 |
| 8 | 707 | 976 | 420 | 825 | 827 | 484 | 584 | 719 | 144 | 408 | 414 | 542 |
| 9 | 0 | 0 | 1 | 499 | 907 | 6 | 890 | 972 | 25 | 948 | 978 | 17 |
| 10 | 146 | 251 | 0 | 841 | 926 | 19 | | | | | | |
| 11 | 37 | 990 | 88 | 77 | 83 | 33 | | | | | | |
| 12 | 830 | 985 | 508 | 508 | 543 | 268 | | | | | | |
| 13 | 3 | 39 | 292 | 56 | 147 | 96 | | | | | | |
| 14 | 385 | 527 | 529 | 22 | 32 | 46 | | | | | | |
| 15 | 33 | 290 | 73 | 10 | 18 | 910 | | | | | | |
| 16 | 94 | 150 | 0 | 364 | 467 | 236 | | | | | | |
| 17 | 12 | 12 | 887 | 9 | 15 | 29 | | | | | | |
| 18 | 2 | 9 | 280 | 958 | 976 | 230 | | | | | | |
| 19 | 124 | 993 | 1 | 6 | 6 | 16 | | | | | | |
| 20 | 0 | 10 | 0 | 30 | 176 | 596 | | | | | | |
| 21 | 278 | 313 | 28 | 752 | 785 | 224 | | | | | | |
| 22 | 0 | 0 | 666 | 874 | 953 | 129 | | | | | | |
| 23 | 5 | 165 | 117 | 23 | 23 | 909 | | | | | | |
| 24 | 4 | 37 | 85 | 580 | 589 | 91 | | | | | | |
| 25 | 5 | 10 | 772 | 266 | 485 | 28 | | | | | | |
| 26 | 0 | 10 | 1 | 302 | 489 | 447 | | | | | | |
| 27 | 246 | 996 | 17 | 114 | 166 | 681 | | | | | | |
| 28 | 40 | 953 | 177 | 171 | 178 | 245 | | | | | | |
| 29 | 576 | 733 | 27 | 11 | 115 | 420 | | | | | | |
| 30 | 46 | 47 | 760 | 32 | 38 | 0 | | | | | | |
| 31 | 639 | 644 | 0 | 897 | 901 | 97 | | | | | | |
| 32 | 60 | 91 | 115 | 513 | 667 | 0 | | | | | | |
| 33 | 26 | 988 | 0 | 232 | 254 | 9 | | | | | | |
| 34 | 897 | 968 | 336 | 5 | 13 | 981 | | | | | | |
| 35 | 455 | 590 | 125 | 668 | 672 | 0 | | | | | | |
| 36 | 12 | 13 | 25 | 71 | 99 | 76 | | | | | | |
| 37 | 68 | 152 | 1 | 985 | 999 | 989 | | | | | | |
| 38 | 61 | 992 | 1 | 590 | 745 | 318 | | | | | | |
| 39 | 89 | 90 | 1 | 940 | 1000 | 2 | | | | | | |
| 40 | 29 | 34 | 113 | 8 | 8 | 440 | | | | | | |
| 41 | 12 | 12 | 218 | 957 | 970 | 923 | | | | | | |
| 42 | 709 | 919 | 0 | 918 | 919 | 109 | | | | | | |
| 43 | 388 | 999 | 626 | 0 | 0 | 2 | | | | | | |
| 44 | 0 | 0 | 187 | 8 | 297 | 16 | | | | | | |
| 45 | 4 | 393 | 27 | 372 | 952 | 8 | | | | | | |
| 46 | 210 | 506 | 10 | 360 | 389 | 0 | | | | | | |
| 47 | 88 | 765 | 0 | 6 | 6 | 832 | | | | | | |
| 48 | 4 | 13 | 8 | 87 | 155 | 125 | | | | | | |
| 49 | 87 | 109 | 397 | 651 | 997 | 33 | | | | | | |

<table>
<tr><th colspan="3">Classification</th><th colspan="2">Taxon sampling</th></tr>
<tr><td colspan="3">Archaeplastida</td><td>Rhodelphis spp.</td><td></td></tr>
<tr><td rowspan="4">Pancryptista</td><td colspan="2"></td><td>Microheliella maris</td><td></td></tr>
<tr><td rowspan="3">Cryptista</td><td></td><td>Palpitomonas bilix</td><td>✓</td></tr>
<tr><td rowspan="2">Cryptomonada</td><td>Cryptophyceae</td><td>✓</td></tr>
<tr><td>Goniomonadea</td><td>✓</td></tr>
</table>

| Replicate # | rs50g Sister relationship between Archaeplastida and Cryptomonada | rs50g Archaeplastida | rs50g Rhodophyta + Cryptista | rs100g Sister relationship between Archaeplastida and Cryptomonada | rs100g Archaeplastida | rs100g Rhodophyta + Cryptista |
|---|---|---|---|---|---|---|
| 0 | 142 | 206 | 8 | 91 | 621 | 24 |
| 1 | 24 | 31 | 90 | 631 | 935 | 564 |
| 2 | 17 | 17 | 11 | 93 | 467 | 793 |
| 3 | 395 | 892 | 71 | 151 | 195 | 408 |
| 4 | 702 | 726 | 110 | 0 | 0 | 532 |
| 5 | 64 | 599 | 650 | 492 | 516 | 181 |
| 6 | 754 | 756 | 0 | 21 | 369 | 64 |
| 7 | 626 | 999 | 801 | 1 | 5 | 496 |
| 8 | 142 | 996 | 328 | 145 | 145 | 667 |
| 9 | 1 | 2 | 9 | 253 | 845 | 60 |
| 10 | 81 | 386 | 1 | 349 | 415 | 11 |
| 11 | 376 | 956 | 276 | 15 | 25 | 24 |
| 12 | 696 | 929 | 601 | 17 | 177 | 38 |
| 13 | 0 | 215 | 248 | 5 | 166 | 564 |
| 14 | 96 | 209 | 196 | 0 | 22 | 217 |
| 15 | 6 | 316 | 201 | 6 | 14 | 977 |
| 16 | 156 | 173 | 0 | 184 | 258 | 650 |
| 17 | 10 | 14 | 847 | 0 | 3 | 409 |
| 18 | 2 | 8 | 900 | 789 | 856 | 666 |
| 19 | 564 | 971 | 6 | 0 | 1 | 110 |
| 20 | 0 | 64 | 81 | 7 | 129 | 886 |
| 21 | 269 | 320 | 206 | 237 | 264 | 593 |
| 22 | 0 | 0 | 866 | 459 | 779 | 493 |
| 23 | 39 | 155 | 64 | 0 | 0 | 969 |
| 24 | 0 | 92 | 62 | 71 | 155 | 786 |
| 25 | 4 | 25 | 810 | 87 | 175 | 546 |
| 26 | 0 | 4 | 32 | 56 | 64 | 711 |
| 27 | 538 | 990 | 83 | 7 | 82 | 809 |
| 28 | 15 | 863 | 238 | 25 | 28 | 891 |
| 29 | 76 | 80 | 37 | 1 | 95 | 453 |
| 30 | 11 | 19 | 525 | 2 | 21 | 2 |
| 31 | 436 | 436 | 0 | 206 | 213 | 660 |
| 32 | 18 | 21 | 793 | 201 | 253 | 10 |
| 33 | 7 | 949 | 6 | 76 | 98 | 69 |
| 34 | 816 | 900 | 287 | 2 | 25 | 952 |
| 35 | 141 | 198 | 137 | 27 | 27 | 14 |
| 36 | 6 | 10 | 157 | 10 | 54 | 292 |
| 37 | 53 | 96 | 6 | 919 | 998 | 940 |
| 38 | 87 | 988 | 0 | 182 | 244 | 236 |
| 39 | 10 | 29 | 59 | 852 | 990 | 44 |
| 40 | 7 | 7 | 284 | 0 | 2 | 435 |
| 41 | 2 | 2 | 218 | 647 | 757 | 759 |
| 42 | 576 | 650 | 0 | 705 | 706 | 189 |
| 43 | 425 | 961 | 904 | 1 | 6 | 83 |
| 44 | 0 | 0 | 134 | 7 | 326 | 78 |
| 45 | 0 | 296 | 187 | 192 | 901 | 127 |
| 46 | 29 | 117 | 87 | 76 | 292 | 15 |
| 47 | 49 | 607 | 0 | 0 | 0 | 995 |
| 48 | 0 | 7 | 0 | 3 | 135 | 836 |
| 49 | 18 | 22 | 318 | 579 | 915 | 93 |

| Replicate # | rs150g Sister relationship between Archaeplastida and Cryptomonada | rs150g Archaeplastida | rs150g Rhodophyta + Cryptista | rs200g Sister relationship between Archaeplastida and Cryptomonada | rs200g Archaeplastida | rs200g Rhodophyta + Cryptista |
|---|---|---|---|---|---|---|
| 0 | 17 | 53 | 942 | 446 | 509 | 491 |
| 1 | 31 | 34 | 565 | 62 | 71 | 624 |
| 2 | 932 | 953 | 45 | 864 | 923 | 72 |
| 3 | 676 | 926 | 74 | 772 | 779 | 218 |
| 4 | 0 | 3 | 994 | 17 | 21 | 512 |
| 5 | 674 | 727 | 243 | 861 | 989 | 11 |
| 6 | 37 | 121 | 878 | 294 | 299 | 694 |
| 7 | 172 | 173 | 717 | 346 | 365 | 628 |
| 8 | 56 | 246 | 606 | 12 | 14 | 909 |
| 9 | 532 | 714 | 223 | 353 | 424 | 574 |

| Classification | | | Taxon sampling | |
|---|---|---|---|---|
| Archaeplastida | | | *Rhodelphis* spp. | |
| Pancryptista | | | *Microheliella maris* | |
| Pancryptista | Cryptista | | *Palpitomonas biilix* | |
| Pancryptista | Cryptista | Cryptomonada | Cryptophyceae | ✓ |
| Pancryptista | Cryptista | Cryptomonada | Goniomonadea | ✓ |

| Replicate # | rs50g<br>Sister relationship between Archaeplastida and Cryptomonada | rs50g<br>Archaeplastida | rs50g<br>Rhodophyta + Cryptista | rs100g<br>Sister relationship between Archaeplastida and Cryptomonada | rs100g<br>Archaeplastida | rs100g<br>Rhodophyta + Cryptista |
|---|---|---|---|---|---|---|
| 0 | 207 | 263 | 11 | 0 | 50 | 0 |
| 1 | 32 | 80 | 88 | 144 | 359 | 730 |
| 2 | 3 | 8 | 1 | 122 | 609 | 834 |
| 3 | 366 | 902 | 109 | 52 | 224 | 57 |
| 4 | 327 | 403 | 69 | 0 | 0 | 16 |
| 5 | 122 | 678 | 18 | 0 | 8 | 0 |
| 6 | 0 | 0 | 15 | 3 | 413 | 263 |
| 7 | 135 | 1000 | 920 | 0 | 7 | 262 |
| 8 | 44 | 965 | 401 | 31 | 32 | 974 |
| 9 | 0 | 1 | 7 | 26 | 373 | 64 |
| 10 | 99 | 746 | 0 | 143 | 209 | 13 |
| 11 | 105 | 984 | 358 | 7 | 24 | 3 |
| 12 | 586 | 891 | 925 | 0 | 295 | 2 |
| 13 | 0 | 106 | 91 | 0 | 1 | 186 |
| 14 | 8 | 825 | 39 | 0 | 0 | 53 |
| 15 | 6 | 648 | 453 | 4 | 48 | 981 |
| 16 | 54 | 60 | 0 | 47 | 68 | 52 |
| 17 | 6 | 6 | 774 | 0 | 1 | 0 |
| 18 | 0 | 55 | 738 | 612 | 805 | 45 |
| 19 | 567 | 984 | 0 | 0 | 5 | 541 |
| 20 | 0 | 11 | 168 | 0 | 148 | 869 |
| 21 | 129 | 184 | 0 | 50 | 681 | 104 |
| 22 | 0 | 0 | 939 | 409 | 947 | 234 |
| 23 | 9 | 67 | 0 | 0 | 1 | 961 |
| 24 | 0 | 18 | 98 | 3 | 30 | 54 |
| 25 | 0 | 97 | 918 | 0 | 11 | 735 |
| 26 | 0 | 17 | 347 | 1 | 19 | 97 |
| 27 | 112 | 998 | 47 | 3 | 182 | 58 |
| 28 | 101 | 733 | 73 | 4 | 7 | 941 |
| 29 | 0 | 0 | 52 | 0 | 357 | 20 |
| 30 | 0 | 0 | 811 | 0 | 15 | 2 |
| 31 | 223 | 228 | 1 | 1 | 2 | 325 |
| 32 | 5 | 16 | 126 | 99 | 196 | 18 |
| 33 | 3 | 635 | 40 | 0 | 0 | 110 |
| 34 | 865 | 944 | 317 | 0 | 74 | 155 |
| 35 | 47 | 92 | 108 | 5 | 5 | 0 |
| 36 | 7 | 25 | 299 | 0 | 0 | 632 |
| 37 | 22 | 39 | 31 | 987 | 998 | 944 |
| 38 | 138 | 981 | 0 | 324 | 514 | 276 |
| 39 | 2 | 8 | 8 | 553 | 980 | 0 |
| 40 | 0 | 0 | 208 | 0 | 0 | 795 |
| 41 | 0 | 10 | 306 | 140 | 390 | 851 |
| 42 | 408 | 517 | 0 | 112 | 363 | 137 |
| 43 | 370 | 962 | 776 | 0 | 12 | 24 |
| 44 | 0 | 0 | 322 | 14 | 352 | 46 |
| 45 | 0 | 181 | 89 | 9 | 325 | 4 |
| 46 | 8 | 371 | 0 | 35 | 114 | 10 |
| 47 | 5 | 463 | 0 | 0 | 18 | 304 |
| 48 | 0 | 43 | 0 | 2 | 56 | 923 |
| 49 | 65 | 134 | 488 | 122 | 972 | 4 |

| Replicate # | rs150g<br>Sister relationship between Archaeplastida and Cryptomonada | rs150g<br>Archaeplastida | rs150g<br>Rhodophyta + Cryptista | rs200g<br>Sister relationship between Archaeplastida and Cryptomonada | rs200g<br>Archaeplastida | rs200g<br>Rhodophyta + Cryptista |
|---|---|---|---|---|---|---|
| 0 | 4 | 76 | 910 | 96 | 399 | 601 |
| 1 | 0 | 0 | 71 | 22 | 89 | 791 |
| 2 | 497 | 628 | 17 | 586 | 973 | 15 |
| 3 | 170 | 392 | 21 | 340 | 358 | 67 |
| 4 | 0 | 0 | 664 | 1 | 1 | 7 |
| 5 | 124 | 190 | 266 | 368 | 570 | 1 |
| 6 | 7 | 187 | 800 | 117 | 129 | 398 |
| 7 | 32 | 33 | 825 | 94 | 97 | 554 |
| 8 | 0 | 612 | 91 | 0 | 1 | 288 |
| 9 | 15 | 55 | 60 | 183 | 278 | 719 |

| Classification | | | Taxon sampling | |
|---|---|---|---|---|
| Archaeplastida | | | *Rhodelphis* spp. | |
| Pancryptista | | | *Microheliella maris* | |
| | Cryptista | | *Palpitomonas bilix* | |
| | | Cryptomonada | Cryptophyceae | |
| | | | Goniomonadea | ✓ |

| Replicate # | rs50g Sister relationship between Archaeplastida and Cryptomonada | rs50g Archaeplastida | rs50g Rhodophyta + Cryptista | rs100g Sister relationship between Archaeplastida and Cryptomonada | rs100g Archaeplastida | rs100g Rhodophyta + Cryptista | rs150g Sister relationship between Archaeplastida and Cryptomonada | rs150g Archaeplastida | rs150g Rhodophyta + Cryptista | rs200g Sister relationship between Archaeplastida and Cryptomonada | rs200g Archaeplastida | rs200g Rhodophyta + Cryptista |
|---|---|---|---|---|---|---|---|---|---|---|---|---|
| 0 | 2 | 214 | 0 | 0 | 803 | 144 | 0 | 59 | 938 | 0 | 23 | 976 |
| 1 | 0 | 66 | 0 | 1 | 110 | 665 | 0 | 312 | 665 | 0 | 55 | 890 |
| 2 | 0 | 0 | 71 | 0 | 139 | 270 | 0 | 85 | 915 | 0 | 397 | 602 |
| 3 | 14 | 881 | 364 | 1 | 175 | 752 | 0 | 595 | 400 | 0 | 38 | 962 |
| 4 | 9 | 165 | 668 | 0 | 0 | 385 | 0 | 45 | 673 | 0 | 46 | 944 |
| 5 | 1 | 772 | 743 | 5 | 417 | 5 | 0 | 19 | 930 | 0 | 351 | 648 |
| 6 | 45 | 46 | 14 | 0 | 256 | 496 | 0 | 53 | 947 | 0 | 567 | 432 |
| 7 | 28 | 803 | 385 | 0 | 16 | 448 | 1 | 272 | 719 | 0 | 557 | 442 |
| 8 | 0 | 981 | 738 | 0 | 268 | 291 | 0 | 10 | 983 | 0 | 115 | 862 |
| 9 | 0 | 30 | 1 | 0 | 133 | 841 | 0 | 341 | 655 | 0 | 189 | 811 |
| 10 | 0 | 936 | 1 | 1 | 309 | 0 | | | | | | |
| 11 | 0 | 32 | 76 | 0 | 141 | 889 | | | | | | |
| 12 | 6 | 635 | 1 | 0 | 60 | 276 | | | | | | |
| 13 | 0 | 305 | 371 | 0 | 457 | 990 | | | | | | |
| 14 | 27 | 190 | 84 | 0 | 344 | 817 | | | | | | |
| 15 | 0 | 108 | 544 | 0 | 13 | 997 | | | | | | |
| 16 | 1 | 582 | 0 | 0 | 355 | 970 | | | | | | |
| 17 | 0 | 0 | 829 | 0 | 90 | 577 | | | | | | |
| 18 | 0 | 53 | 57 | 0 | 116 | 928 | | | | | | |
| 19 | 0 | 964 | 697 | 0 | 97 | 23 | | | | | | |
| 20 | 0 | 538 | 67 | 0 | 83 | 773 | | | | | | |
| 21 | 0 | 139 | 912 | 0 | 5 | 689 | | | | | | |
| 22 | 0 | 0 | 948 | 0 | 165 | 832 | | | | | | |
| 23 | 2 | 34 | 501 | 0 | 1 | 1000 | | | | | | |
| 24 | 0 | 154 | 911 | 0 | 19 | 954 | | | | | | |
| 25 | 0 | 77 | 525 | 0 | 285 | 722 | | | | | | |
| 26 | 0 | 165 | 0 | 0 | 13 | 32 | | | | | | |
| 27 | 8 | 55 | 689 | 0 | 349 | 909 | | | | | | |
| 28 | 0 | 809 | 81 | 0 | 156 | 962 | | | | | | |
| 29 | 0 | 76 | 409 | 0 | 155 | 25 | | | | | | |
| 30 | 0 | 0 | 771 | 0 | 0 | 169 | | | | | | |
| 31 | 6 | 13 | 0 | 6 | 46 | 718 | | | | | | |
| 32 | 0 | 51 | 986 | 1 | 149 | 752 | | | | | | |
| 33 | 0 | 997 | 0 | 0 | 646 | 685 | | | | | | |
| 34 | 0 | 102 | 472 | 0 | 51 | 911 | | | | | | |
| 35 | 0 | 724 | 18 | 0 | 13 | 271 | | | | | | |
| 36 | 0 | 0 | 217 | 0 | 141 | 994 | | | | | | |
| 37 | 3 | 72 | 511 | 1 | 830 | 785 | | | | | | |
| 38 | 0 | 993 | 0 | 0 | 195 | 373 | | | | | | |
| 39 | 0 | 0 | 0 | 6 | 242 | 278 | | | | | | |
| 40 | 3 | 102 | 208 | 0 | 17 | 487 | | | | | | |
| 41 | 0 | 32 | 578 | 0 | 640 | 585 | | | | | | |
| 42 | 1 | 638 | 49 | 0 | 1 | 427 | | | | | | |
| 43 | 39 | 478 | 840 | 0 | 76 | 895 | | | | | | |
| 44 | 0 | 16 | 827 | 0 | 382 | 111 | | | | | | |
| 45 | 0 | 389 | 132 | 0 | 684 | 529 | | | | | | |
| 46 | 1 | 280 | 951 | 0 | 458 | 96 | | | | | | |
| 47 | 0 | 252 | 194 | 0 | 28 | 978 | | | | | | |
| 48 | 0 | 248 | 0 | 0 | 289 | 717 | | | | | | |
| 49 | 1 | 88 | 300 | 0 | 105 | 863 | | | | | | |

| Classification | | | Taxon sampling | |
|---|---|---|---|---|
| Archaeplastida | | | *Rhodelphis* spp. | |
| Pancryptista | | | *Microheliella maris* | |
| | Cryptista | | *Palpitomonas biilix* | |
| | | Cryptomonada | Cryptophyceae, | ✓ |
| | | | Goniomonadea | |
